# Electric signal polymorphism predicts dietary niche partitioning in a weakly electric fish

**DOI:** 10.64898/2026.04.23.720378

**Authors:** Sophie Picq, Rita Gorsuch, Rosella Bills, Lauren Koenig, Nestor Ngoua Aba’a, Franck Nzigou, Hans Kevin Mipounga, Elise C. Knobloch, Ray C. Schmidt, Emilie Parkanzky, M. Eric Benbow, Jason. R. Gallant

## Abstract

Electric organ discharge (EOD) waveform diversity in African elephantfish is often attributed to sexual selection, yet EODs also mediate active electrolocation during prey detection, raising the possibility that natural selection on foraging ecology contributes to waveform divergence. *Paramormyrops kingsleyae* exhibits an intraspecific polymorphism where certain populations emit biphasic EODs whereas other populations emit triphasic waveforms. The genes underlying this polymorphism show signatures of selection; the polymorphism persists despite gene flow and is behaviorally discriminable by the fish themselves. If waveform differences influence prey detection during active electrolocation, biphasic and triphasic fish should consume systematically different prey. We tested this prediction using DNA metabarcoding of gut contents from 186 mormyrids representing 16 species across eight sites in Gabon, employing two independent COI primer sets for cross-validation and pairing dietary data with environmental invertebrate sampling to distinguish active prey preference from passive availability. At the community level in the diverse Balé Creek mormyrid assemblage, species identity was the dominant predictor of diet composition (R² ≈ 24%), consistent with phylogenetic signal in foraging ecology. Within *P. kingsleyae*, waveform type was the strongest independent predictor of dietary composition (R² = 5–6%), explaining variance independently of geographic region, sex, body size, and parasitism status — a result concordant across both primer sets. Dietary differences were driven by prey species turnover rather than differential abundance of shared prey, and prey selectivity analyses confirmed that waveform types differ in which prey they actively prefer, not merely in what is locally available. These findings are consistent with natural selection on foraging ecology contributing to the maintenance of EOD waveform polymorphism, though the sensory mechanisms linking subtle waveform differences to prey detection remain an open question.

## Introduction

Weakly electric elephantfish in the family Mormyroidea are composed of approximately 200 species endemic to freshwater habitats throughout Africa. These fish produce species-specific electric organ discharges (EODs) that serve dual functions: communication with conspecifics (Hopkins, 1981) and active electrolocation of objects in their environment (Von Der Emde, 1999). Because EODs mediate both mate recognition and prey detection, waveform diversity in mormyrids is potentially subject to both sexual selection and natural selection, though the relative contributions of these evolutionary forces remain poorly understood.

The genus *Paramormyrops*, a geographically restricted, rapidly diverged “species flock” endemic to western-central Africa, presents a striking example of speciation among mormyrids. Despite being morphologically similar across species, *Paramormyrops* exhibits remarkable EOD diversity (Fig.1; Arnegard et al., 2010; Arnegard & Hopkins, 2003; Sullivan et al., 2002, 2004).

**Figure 1.**
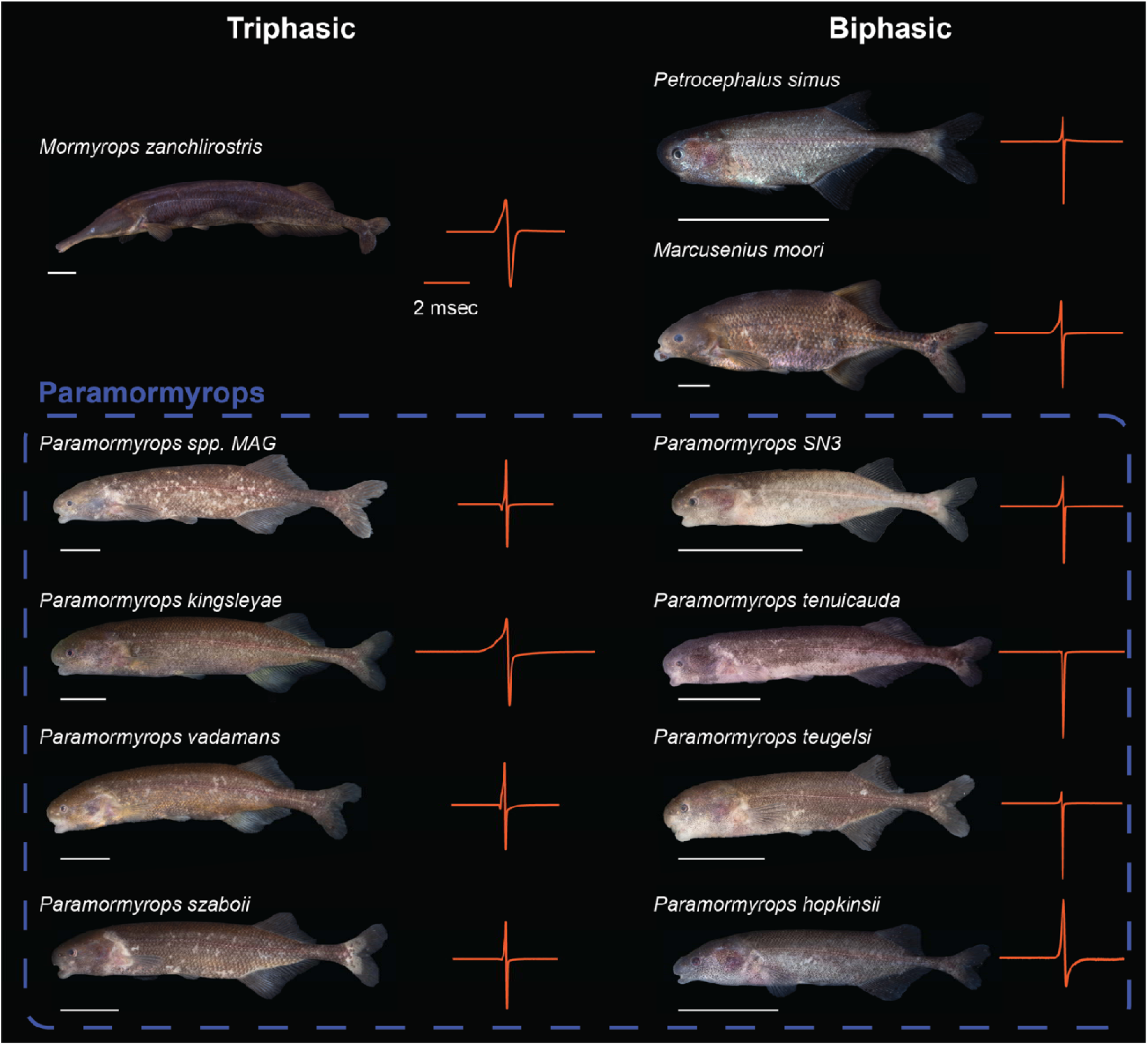
Assemblage of Mormyridae at Balé Creek. Representative images of mormyrid fishes captured at Balé creek together with EOD waveforms (red), which can be categorized as triphasic (left) or biphasic (right). Several *Paramormyrops* (blue dashed line) occur sympatrically with other mormyrid species. White scale bar = 20mm.

Notably, two species — *P. kingsleyae* and *Paramormyrops sp.* ‘MAG’ — exhibit intraspecific polymorphism in EOD waveforms (Arnegard et al., 2005; Gallant et al., 2011; Picq et al., 2020). Where EOD types occur in sympatry, they are genetically indistinguishable, suggesting that EOD polymorphism is maintained despite gene flow (Arnegard et al., 2005; Picq et al., 2020).

*Paramormyrops kingsleyae* is geographically widespread across the lower Guinea Basin (Stiassny et al., 2008) and exhibits substantial geographic variation in EOD waveforms (Gallant et al., 2011; Picq et al., 2020). While this variation was initially attributed to genetic drift under population isolation (Gallant et al., 2011), subsequent work found that patterns of EOD divergence were inconsistent with neutral expectations (Picq et al., 2020), and genomic analyses have revealed signatures of selection at loci associated with EOD divergence (Gallant et al., 2026). These findings raise a fundamental question: what selective pressures are driving EOD divergence?

One potential selective pressure is sexual selection, which has been invoked to explain EOD divergence among *Paramormyrops* (Arnegard et al., 2010) and in *Campylomormyrus* (Feulner et al., 2009a; Tiedemann et al., 2010). However, mormyrids use EODs not only for communication but also for active electrolocation during prey detection (Von Der Emde, 1999), raising an alternative — and not mutually exclusive (Feulner et al., 2009b) — possibility: that natural selection has shaped EOD waveforms through their role in foraging ecology. This alternative is particularly intriguing in *P. kingsleyae*, where BP and TP waveforms have essentially identical power spectra. Moreover, the additional phase distinguishing TP from BP signals contributes negligibly to overall signal amplitude, yet fish can reliably discriminate the two waveforms behaviorally (Picq et al., 2020). How such physically subtle differences could influence foraging is unknown, but if they do, BP and TP fish should consume systematically different prey. Testing this prediction requires dietary data with sufficient taxonomic resolution to detect fine-scale differences in prey composition — a level of resolution that prior approaches have lacked.

Prior dietary work on mormyrids has faced two fundamental limitations. Stable isotope analysis — used to examine trophic ecology in *Paramormyrops* at Balé Creek and Loa Loa Rapids (Arnegard et al., 2010) — places consumers in a bivariate trophic space reflecting general carbon sources and trophic position. However, it cannot resolve prey to finer than broad functional categories and cannot distinguish identical feeding behavior from consumption of functionally equivalent prey in different microhabitats. The first DNA-based diet study of mormyrids (Amen et al., 2024) used hybridization-capture metabarcoding in *Campylomormyrus* to demonstrate that species with different snout morphologies consumed detectably different prey at family and genus level, but could not determine whether differences reflected active preference or differential prey availability. *Paramormyrops* presents a more stringent test than *Campylomormyrus*: its species are morphologically conserved, so dietary differences between waveform types, if they exist, are more likely mediated by sensory differences in electrolocation than by mechanical constraints on feeding.

We address these limitations by applying DNA metabarcoding of gut contents to *P. kingsleyae* and co-occurring mormyrids at Balé Creek — one of the few localities where *P. kingsleyae* is sympatric with a diverse, well-characterized mormyrid assemblage — and at additional sites spanning the geographic range of BP and TP populations. This multi-species community context allows intraspecific dietary variation to be interpreted against the broader dietary landscape of the radiation. We use two independent COI primer sets as built-in cross-validation and pair molecular diet data with environmental sampling of prey availability, enabling direct comparison of what fish eat versus what is available — the critical distinction between prey availability and active preference that prior studies have not addressed. We further developed a local barcode reference database from specimens collected at our study sites, addressing the severe underrepresentation of African freshwater invertebrates in public sequence repositories.

If natural selection on foraging ecology contributes to EOD divergence, a clear prediction follows: fish with different waveforms should eat different things. We begin by characterizing the dietary landscape of the Balé Creek mormyrid assemblage, asking how much dietary overlap exists among sympatric species. Against this community-level backdrop, we ask whether BP versus TP waveform type predicts dietary composition within *P. kingsleyae*, and whether this relationship persists after accounting for geographic variation in prey availability. Resolving these questions bears directly on whether the subtle but measurable difference between BP and TP waveforms has functional consequences for foraging, and whether natural selection on electrolocation could contribute to the maintenance of EOD polymorphism.

## Methods

### Mormyrid Sampling

Mormyrid specimens for this study (n = 186) were collected from 16 species, predominantly focusing on *Paramormyrops kingsleyae*. All specimens were collected in Gabon, West Central Africa, from July - August 2019 from the localities shown in Fig. 2. A comprehensive catalog of these specimens, including metadata and voucher accession numbers for the MSU Museum is provided in Supplementary Data File S1. Specimens were collected using hand nets and electric fish detectors or with hook and line. EODs were recorded within 24 hours of capture in 1-to 5-liter plastic containers filled with water from the collection site. Signals were detected using bipolar chloride-coated silver wire electrodes, amplified (bandwidth = 0.0001-50 kHz) with a differential bioamplifier (CWE, Inc., Ardmore, PA), and digitized at sampling rates ranging from 100 kHz to 1 MHz. Head positivity was plotted upward using a USB-powered A-D converter (National Instruments, Austin, TX). Each recording maintained a 16-bit vertical resolution, and multiple EOD waveforms were collected per specimen. Following EOD recording, specimens were euthanized with an overdose of MS-222. Gut contents were dissected out immediately from euthanized fish and preserved in 95% ethanol. The remainder of each specimen was assigned a unique identification tag and fixed in 10% phosphate-buffered formalin (pH 7.2) for at least two weeks before being transferred to 70% ethanol for long-term storage. Specimens were deposited in the MSU Museum. All methods adhered to protocols approved by MSU Campus Animal Resources and the Institutional Animal Care and Use Committee (IACUC).

**Figure 2.**
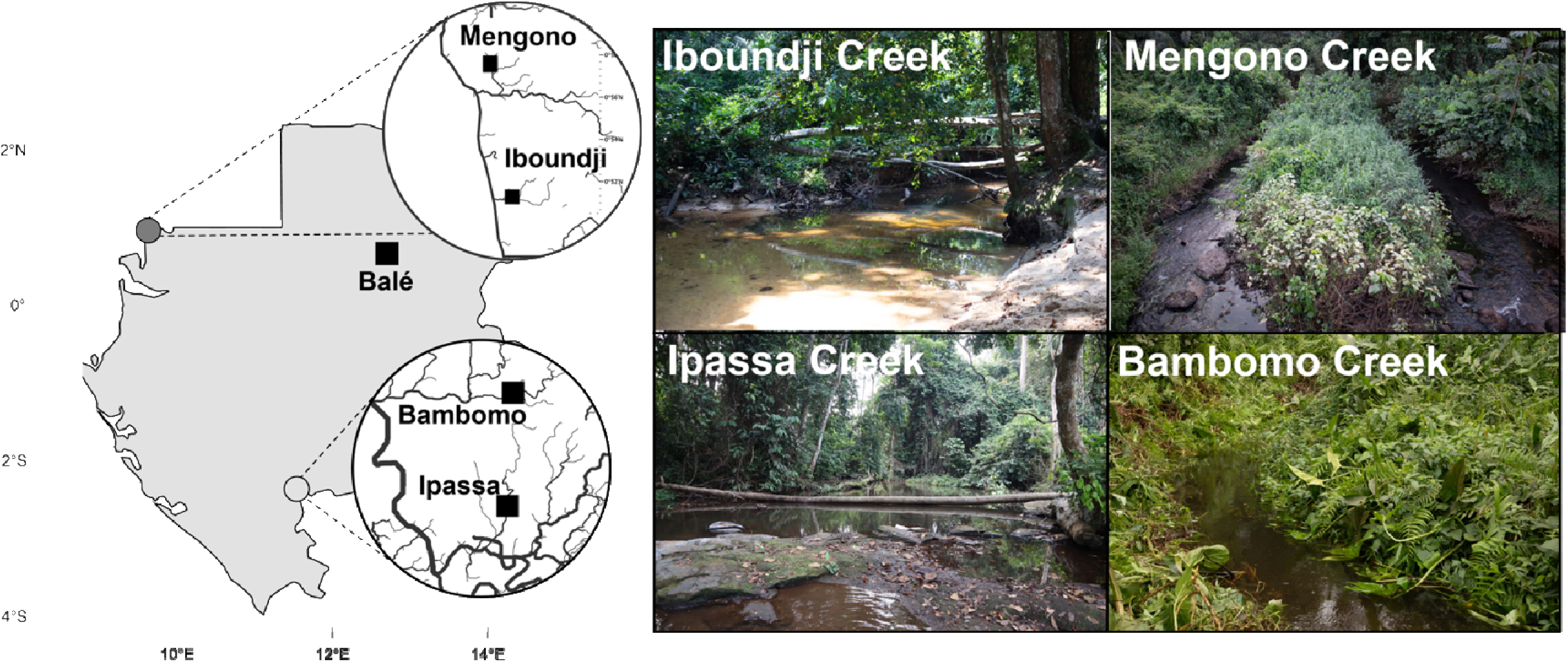
Map of Study Populations and representative photos of habitats.

### Parasite Identification

Morphological identification was performed in consultation with Dr. František Moravec (Institute of Parasitology, Czech Academy of Sciences), a recognized authority on fish nematode taxonomy. Based on photographic examination, the specimens were consistent with either gravid females of family Philometridae or third-stage larvae of Eustrongylides (Dioctophymatidae), both of which can reach similar body sizes and coloration in fish tissue. Discrimination between these taxa was achieved by dissection and examination for the presence of intrauterine larvae, a defining character of viviparous Philometridae.

Genomic DNA was extracted from four individual worm specimens (bc1, bc2,bc3,bc4, extracted from specimens MSU-214, MSU-410, MSU-48, and MSU-412 accordingly) using the Qiagen DNeasy Blood & Tissue Kit following the manufacturer’s protocol. Prior to whole-genome sequencing, conventional PCR-based barcoding was attempted using two primer sets. Universal COI “Folmer” primers (LCO1490/HCO2198; Folmer et al. 1994) and nematode-specific primers JB3/JB4.5 (Derycke et al. 2005) were used to amplify COI from extracted DNA. Amplicons were gel-purified and Sanger sequenced.

Libraries were prepared using the Oxford Nanopore Technologies (ONT) Rapid Barcoding Kit and sequenced on two MinION flow cell for approximately 24 hours each. Basecalling and demultiplexing were performed using standard ONT software, yielding approximately 3,000 reads per specimens bc1 and bc2, and 25,000 reads for specimens bc3 and bc4.

Initial analysis was focused solely on recovering a COX1 barcode sequence for species identification. Flye assembly of nanopore reads from specimen bc4 yielded a COX1-containing contig of approximately 11 kb — far larger than the ∼650 bp barcode fragment anticipated — with the COX1 coding sequence occupying only a small central portion flanked by extensive non-coding sequence of unclear origin. Suspecting either a misassembly artifact or an unusual genomic feature, a literature search was conducted and revealed a highly unusual mitochondrial architecture in *Dioctophyme renale* (Macchiaroli et al., 2025): 11 separate circular minichromosomes, each encoding a single protein-coding gene surrounded by a large non-coding region (NCR). The bc4 assembly was consistent with exactly this structure. This unexpected finding motivated the comprehensive assembly pipeline described below, targeting all 11 mitochondrial minichromosomes in addition to the nuclear rRNA locus.

To identify and assemble sequences of parasitic origin from unassembled nanopore reads, four complementary recruitment strategies were employed. First, reads were screened against a DIAMOND (v2) database of D. renale mitochondrial protein-coding gene (PCG) amino acid sequences using translated alignment (blastx), with invertebrate mitochondrial genetic code (--query-gencode 5), long-read chaining (--long-reads), and sensitive mode. Second, to capture diverged reads that might be missed by the species-specific database, reads were additionally screened against a broader Clade I nematode mitochondrial protein database constructed from 19 reference genomes (Macchiaroli et al. 2025; GenBank accessions NC_071371, NC_056391, NC_008640, NC_008693, NC_008692, NC_025749–NC_025755, NC_002681, NC_018596–NC_018597, NC_017747, NC_017750, NC_028621, NC_001328), using the same DIAMOND parameters with a relaxed e-value cutoff of 1×10⁻³. Third, ribosomal RNA genes were detected using Barrnap v0.9 with the eukaryotic HMM database (--kingdom euk); reads containing 18S, 5.8S, or 28S rRNA annotations were retained (5S rRNA features were excluded as common spurious hits). Fourth, reads were aligned against the BOLD Systems public reference library (downloaded March 27, 2026; 22,992,638 sequences) pre-filtered to Nematoda (61,743 sequences) using a pre-indexed minimap2 v2.28 database (-ax map-ont --secondary=no -f 0.0001). Mitochondrial candidate reads were pooled as the union of the two DIAMOND passes; rRNA candidate reads were pooled as the union of Barrnap and BOLD-mapped reads.

Mitochondrial pool reads were re-queried against the D. renale DIAMOND database in all-hits mode (--max-hsps 0) to assign each read to one or more of the 11 mitochondrial minichromosome targets (mtDNA_01–11, corresponding to ATP6, COX1–3, CYTB, NAD1–6). Per-gene read sets were assembled independently using Flye (--nano-raw --meta --keep-haplotypes --min-overlap 1000 --iterations 3) with target genome sizes of 4 kb per minichromosome and 8 kb for the rRNA locus (18S–ITS1–5.8S–ITS2–28S). To account for non-deterministic Flye behavior at low coverage, assembly was attempted up to three times per target; in cases of complete failure, the deepest available intermediate (post-polishing, post-contigger, or pre-graph consensus) was rescued as the final assembly. Assemblies were subsequently polished with Medaka (medaka_consensus; model r941_min_sup_g507) to correct residual homopolymer errors.

Assemblies for minichromosomes mtDNA_01–10 were validated against concatemerization artifacts using three independent tests: (A) gene copy number, assessed by DIAMOND blastx against the D. renale protein database with all-hits mode; (B) window self-alignment at published minichromosome-size offsets using minimap2, where concatemers would produce >90% inter-window identity; and (C) internal repeat detection within the non-coding region (NCR) upstream of the annotated gene using minimap2 all-versus-all alignment. NAD6 (mtDNA_11) consistently yielded insufficient reads for assembly; an overlap graph and gene-hit inventory were generated to document recoverability.

Mitochondrial minichromosomes (mtDNA_01–10) were annotated with MITOS2 (runmitos.py; invertebrate mitochondrial genetic code 5; RefSeq89 Metazoa reference). The assembled rRNA locus was identified by BLASTn against the NCBI nucleotide (nt) database (e-value ≤ 1×10⁻¹_, up to 10 target sequences), either locally or via remote NCBI query.

### Invertebrate Sampling, Identification, and Barcode Generation

Benthic macroinvertebrates were sampled across aquatic microhabitats (substrate types and both submerged and emergent vegetation) at each river, focusing on areas occupied by mormyrids. Standardized samples were collected using a 500 µm mesh D-frame net by inserting the net 2–3 cm into the sediment facing upstream and disturbing the substrate to dislodge organisms. Floating vegetation was sampled by agitating plants and sweeping the net through them. Plant detritus (e.g., dead leaves and branches) was collected, rinsed into a sieve, and examined individually for macroinvertebrates. All samples were processed at a field laboratory using a series of sieves. Macroinvertebrates were sorted visually by the field team and preserved in 96–100% molecular grade ethanol.

Invertebrates were identified to family (insects) or class/order (non-insect invertebrates) level using Merritt et al. (2008) or Thorp & Covich (2010) at Michigan State University. COI barcodes were generated from identified macroinvertebrate specimens following Knobloch & Schmidt (2022) and Srivathsan et al. (2021). Briefly, ethanol-preserved specimens were rinsed to remove residual ethanol prior to extraction. DNA was extracted non-destructively using QuickExtract (Lucigen) in 96-well format (or from a leg for large specimens), and extracts were diluted prior to amplification; specimens were retained as vouchers. The COI barcode region was amplified using a tagged-primer, combinatorial indexing strategy (12 forward × 24 reverse tags) to enable multiplexing of individual specimens. PCR products were screened on agarose gels to verify amplification, pooled in equal volumes, and purified using AMPure XP magnetic beads to remove primers and short fragments. Purified pools were quantified by fluorometry.

Sequencing libraries were prepared using the Oxford Nanopore ligation kit (SQK-LSK109) following manufacturer guidance and sequenced on Flongle flow cells using a MinION device with MinKNOW basecalling output as FASTQ. Reads were demultiplexed and specimen-level consensus COI barcodes were generated using ONTbarcoder (Srivathsan et al., 2021) with default parameters, using a mapping file linking specimen IDs to tag combinations.

### Stomach Contents: DNA Extraction, PCR Amplification, and Amplicon Sequencing

DNA was extracted from individual stomach-content samples using the QIAamp Fast DNA Stool Mini Kit (QIAGEN). Stomach contents were pelleted by centrifugation (3,700 rpm), the supernatant ethanol was removed, and the pellet was rinsed with ddH₂O. Up to 250 mg of stomach-content material was transferred to lysis tubes and combined with 600–1,000 µL InhibitEX Buffer (volume scaled to starting material). Samples were homogenized by bead beating at 5.0 m s⁻¹ for 4 min (four 1-min cycles with 10 s dwell between cycles). DNA extraction then followed the manufacturer’s protocol, and DNA was eluted in ATE buffer and stored at −20 °C until PCR.

We amplified portions of the mitochondrial cytochrome c oxidase subunit I (Leray et al., 2013) using two independent primer sets (Primer Set 1: n = 166; Primer Set 2: n = 121; primer sequences and annealing temperatures in Table 1). A subset of samples (n = 115) was amplified with both primer sets to enable cross-validation of downstream analyses. A full mapping of samples to primer sets is provided as Supplemental Files 2 and 3. Each 25 µL PCR contained 2.5 µL 10× TaqBuffer II, 2.0 µL MgCl₂, 0.5 µL 10 mM dNTPs, 1.5 µL each of 10 µM forward and reverse primers, 1.0 µL BSA (20 mg mL⁻¹), 0.2 µL Taq DNA polymerase (5 U µL⁻¹), and 2.0 µL template DNA, with nuclease-free water to volume. Thermocycling conditions were: denaturation at 95 °C for 10 min; 35 cycles of denaturation at 95 °C for 15s, annealing at 50–54 °C for 30 s (Table 1), and 72 °C for 60 s; followed by a final extension at 72 °C for 5 min. Three PCR replicates were performed per stomach extraction.

**Table 1.**
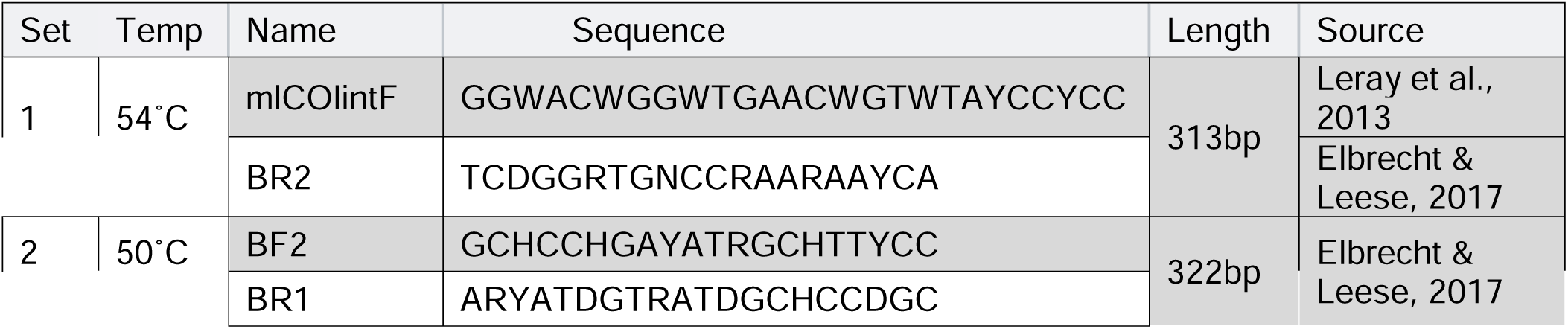
Annealing temperatures, primer names, and sequences used in this study.

Indexed Illumina libraries were generated at the MSU RTSF Genomics Core using a secondary PCR to add dual indices and Illumina-compatible adapters via primers targeting the CS1/CS2 ends of the primary amplicons. Libraries were batch-normalized (Invitrogen SequalPrep DNA Normalization plates) and pooled. Pool quality and concentration were assessed using Qubit dsDNA HS, Agilent 4200 TapeStation (HS DNA1000), and KAPA Illumina Library Quantification qPCR assays. The final pool was sequenced on an Illumina MiSeq using a v2 Nano flow cell with 2×250 bp paired-end chemistry (MiSeq v2, 500-cycle kit). Custom sequencing and index primers complementary to the Fluidigm CS1/CS2 oligomers were added to the appropriate reagent cartridge wells. Base calling was performed with Illumina Real Time Analysis (RTA) v1.18.54, and reads were demultiplexed and converted to FASTQ format using bcl2fastq v2.19.1.

### Bioinformatic Analysis

Base called FASTQ files for each amplicon (Primer Set 1 and Primer Set 2) were subjected to various downstream processing steps separately using the QIIME2 (Bolyen et al., 2019) data platform. First, FASTQ files were imported to QIIME2, then subjected to adapter and primer trimming using Cutadapt (Martin, 2011). Sequences were then denoised using DADA2 (Callahan et al., 2016) retaining only the first 224bp for forward reads and 230bp for reverse reads (Primer Set 1) or the first 230bp for forward reads and 221bp for reverse reads (Primer Set 2). Next, sequences were clustered into representative sequences at 98.5% similarity using VSEARCH (Rognes et al., 2016).

Resulting features present in our dataset were classified based on a custom reference database. Starting with the 5-6-2022 release of the COInr database (Meglécz, 2022), we added the 429 barcodes generated from our invertebrate sampling using MkCOInr (Meglécz, 2023).

Sequence features were classified using two methods: first representative sequences with 80% identity and 94% match to existing barcodes were identified using VSEARCH, and second, we trained a naïve Bayesian classifier using QIIME2 to our custom database and then classified sequences using feature-classifier classify-sklearn with a confidence threshold of 30% (Bokulich et al., 2018; Pedregosa et al., 2011). We merged the two classifications following the methods of O’Rourke et al. (2021) by prioritizing matches from VSEARCH at the family level, then retaining classifications made by the naïve Bayesian classifier at least to the family level.

### Statistical Analysis

All statistical analyses were performed in R version 4.4.0 (Team, 2024). Key packages included *phyloseq* v1.50.0 for microbiome data handling (McMurdie & Holmes, 2013), *vegan* v2.7-2 for community ecology analyses (Oksanen et al., 2022), *qiime2R* v0.99.6 for importing QIIME2 artifacts (Bisanz, 2018), *pairwiseAdonis* v0.4.1 for pairwise PERMANOVA (Martinez Arbizu, 2020), *spaa* v0.2.5 for niche overlap calculations (Zhang, 2016), *car* for Type II ANOVA (Fox & Weisberg, 2019), *FSA* for Dunn’s post-hoc tests (Ogle et al., 2023), *effsize* for effect size calculations, *pheatmap* for heatmap visualization, *ape* for phylogenetic analysis (Paradis & Schliep, 2019), and *cowplot* for plot composition (Wilke, 2024). Visualizations were created using *ggplot2* v3.5.2 (Wickham, 2016) with multi-panel figures assembled using *patchwork* (Pedersen, 2024). Statistical significance was assessed at α = 0.05 throughout, with Benjamini-Hochberg correction applied for multiple comparisons unless otherwise noted. All analysis scripts are available at https://github.com/msuefishlab/gut_contents_2025.

### Host Sequence Filtering and Rarefaction

Prior to diversity analyses, amplicon sequence variants (ASVs) classified as Mormyridae (host DNA) were removed from the feature tables. This filtering step was performed before rarefaction to ensure that differences in host DNA amplification between samples did not bias estimates of prey diversity. Following host removal, samples were rarefied to 3,000 sequences per sample to standardize sequencing depth across all specimens. Samples with fewer than 3,000 prey sequences after host removal were excluded from downstream analyses. For community-level analyses at Balé Creek and within *Paramormyrops kingsleyae*, rare taxa present in fewer than two samples or with fewer than ten total reads were removed to reduce noise from spurious sequences. To validate the robustness of our findings across primer sets, all statistical analyses were conducted independently on data from Primer Set 1 and Primer Set 2 (Table 1), with concordance assessed through correlation analyses and Mantel tests on samples successfully amplified by both primer sets.

### Alpha Diversity

Dietary diversity within individual fish species was quantified using the Shannon diversity index, calculated from rarefied ASV counts using the estimate_richness function in the phyloseq R package. Normality of Shannon diversity values was assessed using Shapiro-Wilk tests; when assumptions of normality were met (P > 0.05), parametric tests were used, and when data were non-normally distributed, non-parametric alternatives were applied. For the Balé Creek community analysis, Shannon diversity was compared across species using one-way ANOVA or Kruskal-Wallis tests based on normality. For *P. kingsleyae* analyses, two-way ANOVA was employed with region (North vs. South) and waveform type (BP vs. TP) as fixed factors, including tests for their interaction. Additional alpha diversity comparisons examined sex (using ANCOVA with body size as a covariate to control for sexual size dimorphism), body size (linear regression with standard length as a continuous predictor), and parasitism (Kruskal-Wallis or ANCOVA controlling for body size). To assess concordance between primer sets, Pearson correlation coefficients were calculated for Shannon diversity values from samples successfully amplified by both primer sets.

### Beta Diversity and Comprehensive PERMANOVA Framework

Dietary composition differences among groups were assessed using Bray-Curtis dissimilarity calculated from rarefied ASV abundances with the vegdist function in the vegan R package. Dissimilarity matrices were visualized using Principal Coordinates Analysis (PCoA) implemented with the cmdscale function. Statistical significance of compositional differences was tested using permutational multivariate analysis of variance (PERMANOVA) with 999 permutations, implemented using the adonis2 function in vegan.

Rather than testing each predictor in isolation, we adopted a comprehensive additive PERMANOVA framework that simultaneously fit all relevant predictors, allowing each factor’s contribution to be assessed while controlling for the others. For the Balé Creek mormyrid community, species with fewer than three sampled individuals were excluded to ensure stable estimates, and PERMANOVA was performed with species identity, waveform type, sex, and body size as simultaneous predictors using marginal (Type III) tests to assess independent contributions. Pairwise PERMANOVA comparisons among species pairs were conducted using the pairwiseAdonis R package with Benjamini-Hochberg correction. For *P. kingsleyae*, a comprehensive PERMANOVA model included waveform type, geographic region, sex, body size (standard length), and parasitism status as simultaneous predictors, again with marginal tests to isolate the independent contribution of each factor after controlling for all others.

Robustness to term ordering was confirmed by running sequential PERMANOVA tests with both orderings of waveform and region (by = “terms”). Homogeneity of multivariate dispersions was assessed for each factor using the betadisper function in vegan, with significance tested by permutation.

To assess concordance between primer sets, Mantel tests were performed on Bray-Curtis distance matrices from samples amplified by both primer sets, using the mantel function in vegan with 999 permutations and Pearson correlation.

### Variance Partitioning

To decompose variation in diet composition into independent and shared components, we performed variance partitioning using the varpart function in vegan applied to Bray-Curtis distance matrices. For Balé Creek, three-way variance partitioning separated the unique contributions of species identity, waveform type, and individual-level factors (sex + body size). For *P. kingsleyae*, two- and three-way variance partitioning decomposed dietary variation into pure waveform effects, pure geographic effects, pure individual-level effects (sex, body size, parasitism), and their shared components. Adjusted R² values were used to account for the number of parameters in each predictor set.

### Prey Taxa Driving Dietary Differences

Similarity percentage analysis (SIMPER; Clarke, 1993) was used to identify which prey taxa contributed most to between-group dissimilarity, implemented using the simper function in vegan with 999 permutations. SIMPER was performed at the ASV level with results aggregated to the family level. This analysis was applied across all significant predictors identified in the comprehensive PERMANOVA models: among Balé Creek species, between waveform types, regions, sexes, body size categories, and parasitism groups within *P. kingsleyae*. For prey families contributing ≥5% to overall dissimilarity, we conducted analyses to partition Bray-Curtis dissimilarity into turnover (prey replacement) and abundance (differential abundance of shared prey) components. For each contributing family, we calculated Jaccard similarity of ASV composition between groups, tested abundance differences in shared ASVs using Wilcoxon rank-sum tests with false discovery rate (FDR) correction, and partitioned Bray-Curtis dissimilarity into turnover and abundance components using presence-absence (Jaccard) and abundance-based (Bray-Curtis) metrics.

### Prey Selectivity Analysis

To assess whether mormyrids feed selectively relative to prey availability, we compared gut content composition with environmental prey availability data from D-frame net collections at the same sites. Selectivity was calculated at the individual fish level: for each specimen, gut content ASV counts were aggregated to family level and compared to the family-level environmental prey assemblage at its collection locality. Selectivity was quantified using Ivlev’s Electivity Index: E = (r − p) / (r + p), where r is the proportion of a prey taxon in the diet and p is its proportion in the environment (Ivlev, 1961). Values range from −1 (complete avoidance) to +1 (maximum preference), with 0 indicating random feeding proportional to availability. Prey families with |E| > 0.2 were classified as preferred or avoided. To test whether selectivity patterns differed between waveform types, individual selectivity profiles were converted to wide-format matrices and PERMANOVA was performed on Euclidean distance matrices of Ivlev’s E values with waveform and region as factors (marginal tests). Overall selectivity intensity was compared between waveform types using mean absolute selectivity (|E|) per individual fish, tested with Wilcoxon rank-sum tests.

## Results

### Overall Results

DNA metabarcoding of gut contents was performed on 186 individual mormyrid fish representing 16 species collected from eight sites across three geographic regions in Gabon. Of these, 119 were *Paramormyrops kingsleyae*—the focal species for waveform-diet analyses.

Two independent COI primer sets were employed for cross-validation: Primer Set 1 (mlCOIintF/BR2) was used to sequence 183 fish, while Primer Set 2 (BF2/BR1) was used to sequence 146 fish. Critically, 146 fish were sequenced with both primer sets to enable direct cross-validation of dietary patterns within individuals. The bioinformatics pipeline included sequential quality filtering steps that differentially affected sample retention between primer sets: adapter trimming, DADA2 denoising and chimera removal, family-level taxonomic filtering (retaining only ASVs assigned to at least Family rank), host DNA removal (filtering Chordata sequences), and rarefaction to 3,000 sequences per sample. After all quality control steps, PS1 retained 179 samples with 4,949 unique prey ASVs (amplicon sequence variants) representing 8.04 million reads, while PS2 retained 142 samples with 3,825 unique ASVs representing 4.34 million reads.

### Parasite Identification

Morphological Identification No intrauterine larvae were detected in either specimen, excluding Philometridae and providing morphological support for identification as Eustrongylides sp. larvae, consistent with the molecular evidence described below.

Conventional Barcoding Amplification with Folmer Primers yielded sequences that BLASTn identified exclusively as *Paramormyrops kingsleyae*, consistent with predominant host DNA in the extracts and corroborating the host-derived 28S rRNA sequence subsequently recovered by ONT sequencing (see below). Amplification with nematode-specific primers JB3/JB4.5 failed to produce reliable bands, precluding Sanger-based nematode identification. motivating the whole-genome ONT sequencing approach described below.

From 63,549 total nanopore reads, multi-strategy recruitment identified 999 candidate reads of parasitic origin. DIAMOND-based translated alignment against *D. renale* mitochondrial protein-coding genes and a broader Clade I nematode mitochondrial protein database together recruited 675 reads to the mitochondrial pool. An additional 332 reads were recruited to the rRNA pool via Barrnap rRNA detection (98 reads bearing 18S, 5.8S, or 28S annotations) supplemented by minimap2 alignment against the BOLD Nematoda reference library.

Flye assembly of the mitochondrial read pool produced contiguous assemblies for 10 of the 11 *D. renale*-type mitochondrial minichromosomes (mtDNA_01 through mtDNA_10; 121,950 bp total), spanning all annotated protein-coding genes except NAD6. Individual minichromosome lengths ranged from 4,004 bp (mtDNA_05, CYTB) to 22,044 bp (mtDNA_06, NAD1), consistently exceeding the corresponding published *D. renale* minichromosome sizes (3,834–4,640 bp; Macchiaroli et al. 2025) by 1.3-to 5.7-fold. Assembly integrity validation — using gene copy number assessment, window self-alignment at published-size offsets, and NCR internal-repeat detection — confirmed these expansions represent genuine non-coding region (NCR) enlargements rather than assembly or concatemerization artifacts. MITOS2 annotation confirmed the expected single protein-coding gene per minichromosome in all assemblies where annotation succeeded; mtDNA_08 (NAD3+NAD4L) was the most feature-rich, yielding 20 annotated elements including 10 tRNA genes. Assembly of mtDNA_11 (NAD6) was not achieved: only 9 reads were recruited, of which 2 produced weak DIAMOND hits (48.4% and 50.0% identity), and the overlap graph showed insufficient coverage for contig formation. Targeted re-sequencing of this minichromosome is recommended. This mitochondrial assembly is included as a Supplemental File 4.

Assembly of the rRNA read pool produced three contiguous units totaling 13,218 bp (5,684 bp + 4,088 bp + 3,446 bp). The largest unit (5,684 bp) contained a complete 18S–ITS1–5.8S–ITS2–28S ribosomal DNA structure as confirmed by ITSx; BLASTn of the 18S (SSU) sequence against the NCBI nucleotide database returned 98.5% identity to *Dioctophyme renale* strain CIFRI.DR-2 (OQ933019.1) and 99.2% identity to *Eustrongylides* sp. IP1 (LC902677.1). A second partial unit (3,446 bp) yielded 92.1% SSU identity to *D. renale* strains and 91.8% to *Eustrongylides ignotus* (MK340916.1). The ITS region (ITS1 + 5.8S + ITS2 = 765 bp; 329 + 155 + 281 bp) was extracted and submitted to GenBank with accession PZ255420.

The mtDNA_02 minichromosome (COX1; 11,466 bp, circular) was validated against 20 published *Eustrongylides* COX1 reference sequences, all of which mapped at MAPQ=60. A 1,206 bp COX1 coding sequence was extracted across two exons (795 bp and 411 bp) separated by a 13 bp region confirmed by 19 spanning reference sequences and mean 28.5× ONT read coverage. This COX1 was submitted to GenBank with accession PZ259218.

Together, the morphological evidence (absence of intrauterine larvae), the assembled rRNA locus (99.2% 18S identity to *Eustrongylides* sp. IP1), and the COX1 barcode alignment support identification of the endoparasitic specimens as *Eustrongylides* sp. (Nematoda: Dioctophymatida: Dioctophymatidae).

### Bioinformatics Pipeline Performance

Family-level taxonomic filtering, which retained only ASVs with both Order and Family assignments, removed different proportions of sequences between primer sets. PS1 lost 10.0% of ASVs representing 2.0% of total reads (161,262 reads), while PS2 lost 50.9% of ASVs representing 9.1% of total reads (436,397 reads). This five-fold difference in ASV filtering loss (Chi-square test: χ² = 353,513, p < 0.0001) reflects differences in reference database coverage for the prey taxa preferentially amplified by each primer set rather than differential data quality. Critically, 99.3% (PS1) and 99.9% (PS2) of filtered ASVs were completely unassigned sequences lacking even Kingdom-level taxonomy—likely non-target environmental DNA or highly divergent sequences without database representation. Only 0.5-0.1% of filtered ASVs had Order assignments without Family assignments, representing primarily Ephemeroptera (mayfly) sequences that lacked species-level resolution in the reference database. Deeper taxonomic resolution varied modestly between primer sets: PS1 assigned 55.0% of ASVs to genus and 35.6% to species, while PS2 assigned 54.7% to genus and 29.9% to species.

### Reference Database Performance and Local Barcode Contribution

Taxonomy assignments drew from two complementary reference sources: the public COInr Metazoa database and 429 locally-generated COI barcodes from macroinvertebrates collected at the study sites. The contribution of local barcodes varied substantially between primer sets, reflecting genuine differences in which prey taxa each primer set amplifies most efficiently (Fig. S1). For PS1, local barcodes contributed to 58.3% of ASVs but only 34.6% of total reads (3,678 reads), while for PS2, local barcodes contributed to 33.6% of ASVs but 75.6% of total reads (436,302 reads). This cross-primer variation in local barcode contribution was statistically significant for PS2 (Chi-square test: χ² = 12.45, p < 0.001) but not for PS1 (χ² = 0.67, p = 0.414), demonstrating that primer-specific amplification biases interact with reference database composition.

Local barcodes proved particularly valuable for regionally-dominant prey families that are underrepresented in global databases. For PS2, local barcodes provided the sole reference coverage for Heptageniidae (115,211 reads across 2 ASVs) and Caenidae (2,795 reads, 1 ASV) and contributed 88.3% of assignments within Chironomidae—the most abundant prey family overall (358,884 total reads across 78 ASVs). For PS1, local barcodes provided exclusive coverage for Chironomidae (2,402 reads, 7 ASVs), Baetidae (932 reads, 1 ASV), and Caenidae (344 reads, 3 ASVs). Across both primer sets, four prey families had coverage exclusively from local barcodes, and an additional two families showed >50% local contribution, demonstrating that investment in regional reference barcode generation filled critical gaps in global databases for Central African aquatic invertebrate diversity.

### Cross-Primer Validation and Compositional Patterns

The two primer sets amplified partially overlapping but compositionally distinct prey communities, consistent with well-documented primer-specific taxonomic biases in COI metabarcoding (Fig. S2). PS1 recovered Chironomidae (Diptera) as the dominant prey family (34.3% of reads, 1.37 million reads), followed by Megascolecidae (Annelida) (6.8%, 273,016 reads), Hydroptilidae (Coleoptera) (5.2%, 208,023 reads), Carabidae (Coleoptera) (4.0%, 159,324 reads), and Sciaridae (Diptera) (3.8%, 151,363 reads). PS2 also recovered Chironomidae as dominant (36.1%, 1.54 million reads), but subsequent families differed: Isotomidae (Collembola) (6.4%, 271,025 reads), and three families of Ephemeroptera: Heptageniidae (5.3%, 223,992 reads), Caenidae (3.9%, 166,252 reads), and Leptophlebiidae (3.6%, 152,105 reads). These rank-order differences reflect known primer binding site preferences, with PS1 showing stronger amplification of certain Coleoptera and microcaddisflies (Trichoptera: Hydroptilidae), while PS2 more efficiently detected mayfly diversity.

The most striking compositional difference involved Hydroptilidae, which represented 5.2% of PS1 reads but only 0.1% of PS2 reads—a 43-fold difference. This family-specific amplification bias likely reflects differential primer binding efficiency to Hydroptilidae COI sequences rather than taxonomic misassignment, as both primer sets achieved 100% family-level assignment rates for recovered sequences. Such primer-specific biases are well-documented in metabarcoding studies and underscore the value of multi-primer approaches for comprehensive prey detection across diverse invertebrate orders.

Despite these compositional differences in specific prey families, both primer sets revealed consistent biological patterns in downstream analyses of dietary variation among mormyrid species and between *P. kingsleyae* waveform types (see subsequent sections). When directional effects were detected (e.g., waveform differences in diet composition), the direction and magnitude of effects were concordant between primer sets, even when specific prey families driving those differences varied.

### Balé Creek

To assess dietary variation among co-occurring mormyrid species while controlling for geographic effects, we analyzed the multi-species mormyrid community at Balé Creek. After complete-case filtering (Table S1), 50 (PS1) and 46 (PS2) individuals representing 9 species were retained for analysis, with five well-sampled species (≥6 individuals per primer set): *Paramormyrops* spp. ‘SZA’, *Marcusenius moori*, *Paramormyrops kingsleyae*, *Petrocephalus simus*, and *Mormyrops zanchlirostris*. Mormyrids of Balé Creek primarily consumed chironomids (>30% of diet composition, Fig. 4A), though the remaining prey consisted of highly diverse taxa including Hydroptilidae, Simuliidae (Diptera), Dolichopodidae (Diptera), and Ephemerellidae (Ephemeroptera) (Table 2). Alpha diversity, measured using Shannon diversity varied significantly among species (PS1: F_8,41_ = 12.56, *P* < 0.001; PS2: F_8,37_ = 2.69, *P* = 0.020; Fig. 4B). Species such as *Mormyrops zanchlirostris* exhibited low diversity indices consistent with dietary specialization, whereas most *Paramormyrops* species showed higher and more similar diversity values.

**Figure 3.**
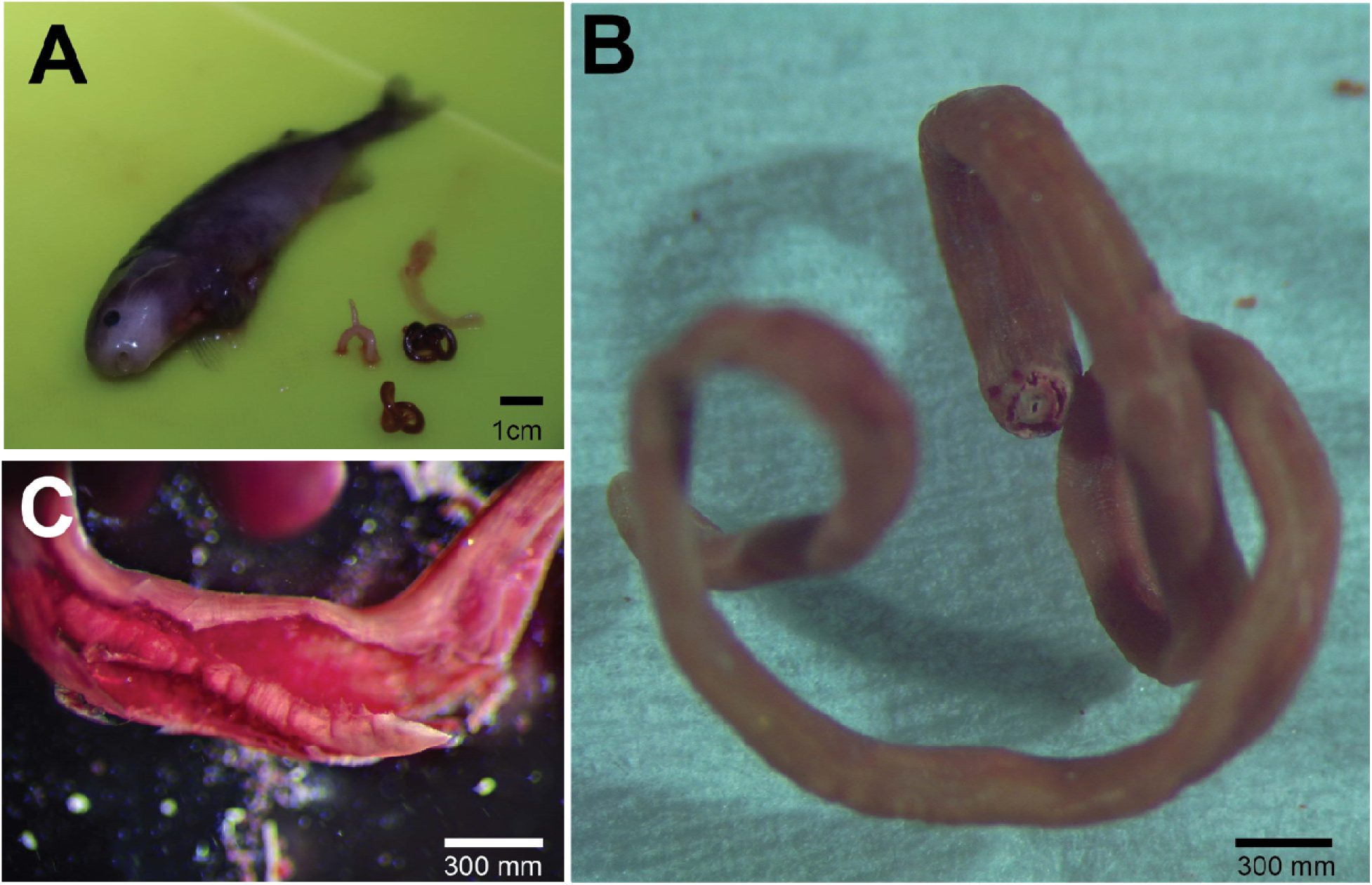
Morphological identification of *Eustrongylides* sp. larvae from *Paramormyrops kingsleyae*. (A) Dissected *P. kingsleyae* specimen (with extracted nematode parasites visible to the right. (B) Stereomicroscopic view of an intact nematode specimen showing the characteristic coiled posture, reddish-pink coloration, and robust body typical of *Eustrongylides*. The blunt anterior end is visible at center. (C) Transverse dissection of nematode body wall revealing internal contents. The absence of intrauterine larvae excludes family Philometridae (viviparous), supporting identification as *Eustrongylides* sp. larvae, consistent with the molecular data.

**Figure 4.**
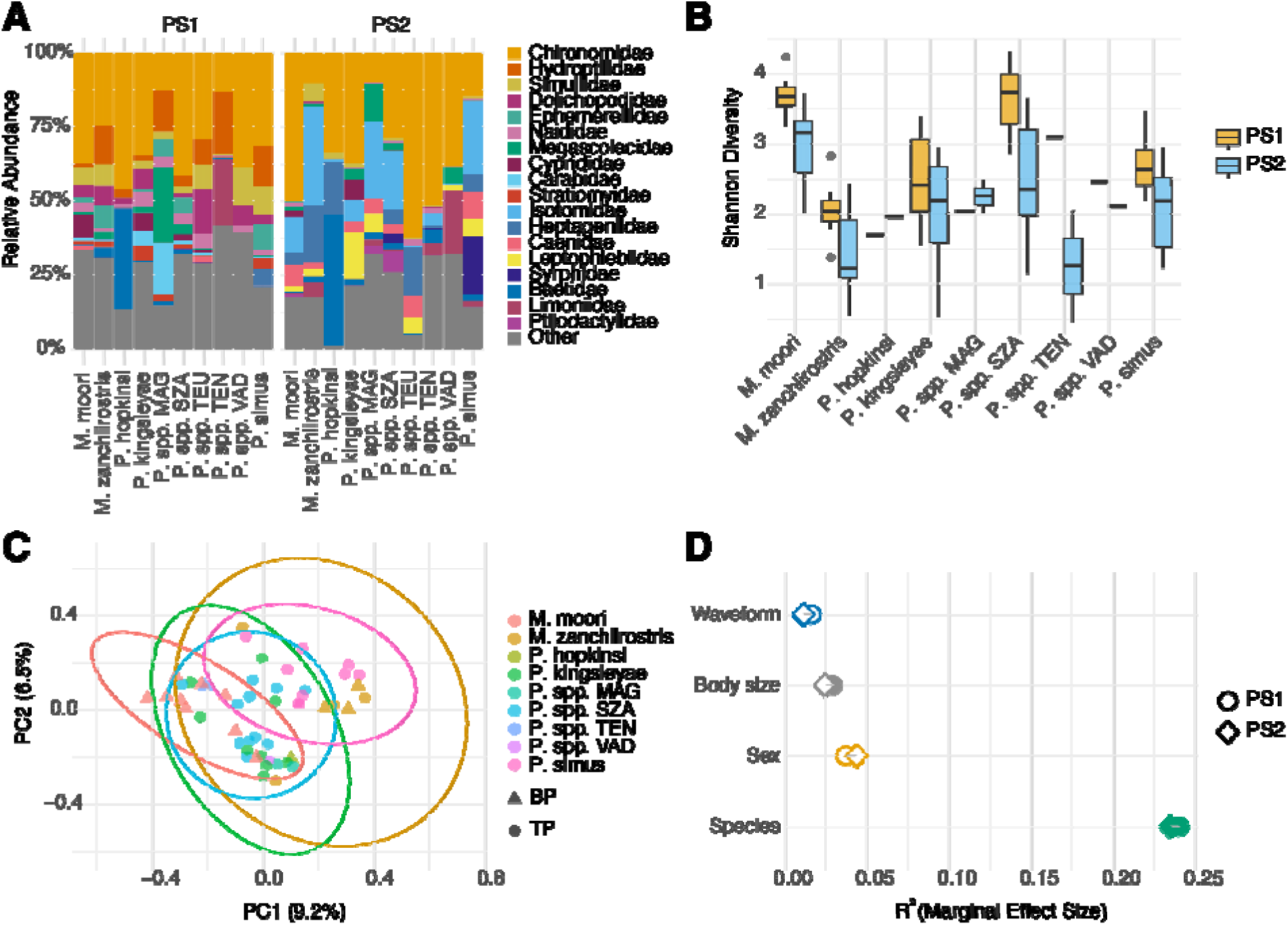
Dietary Diversity in a Community of Balé Creek Mormyrids. A. Composition barplot of the top 20 prey taxa for Primerset 1 and Primerset 2 for each species. B. Mean Shannon diversity indices for each species, colored by primerset. C. PCoA of Bray-Curtis dissimilarity among species for Primerset 1. 95% centroids for each species are shown, biphasic (BP) individuals are shown as triangles, triphasic (TP) individuals are shown as circles. D. Adjusted marginal effect sizes for variance partitioning test of waveform, body size, sex, and species. Filled symbols are significant (p<0.05), empty symbols are not significant (p>0.05).

**Table 2.**
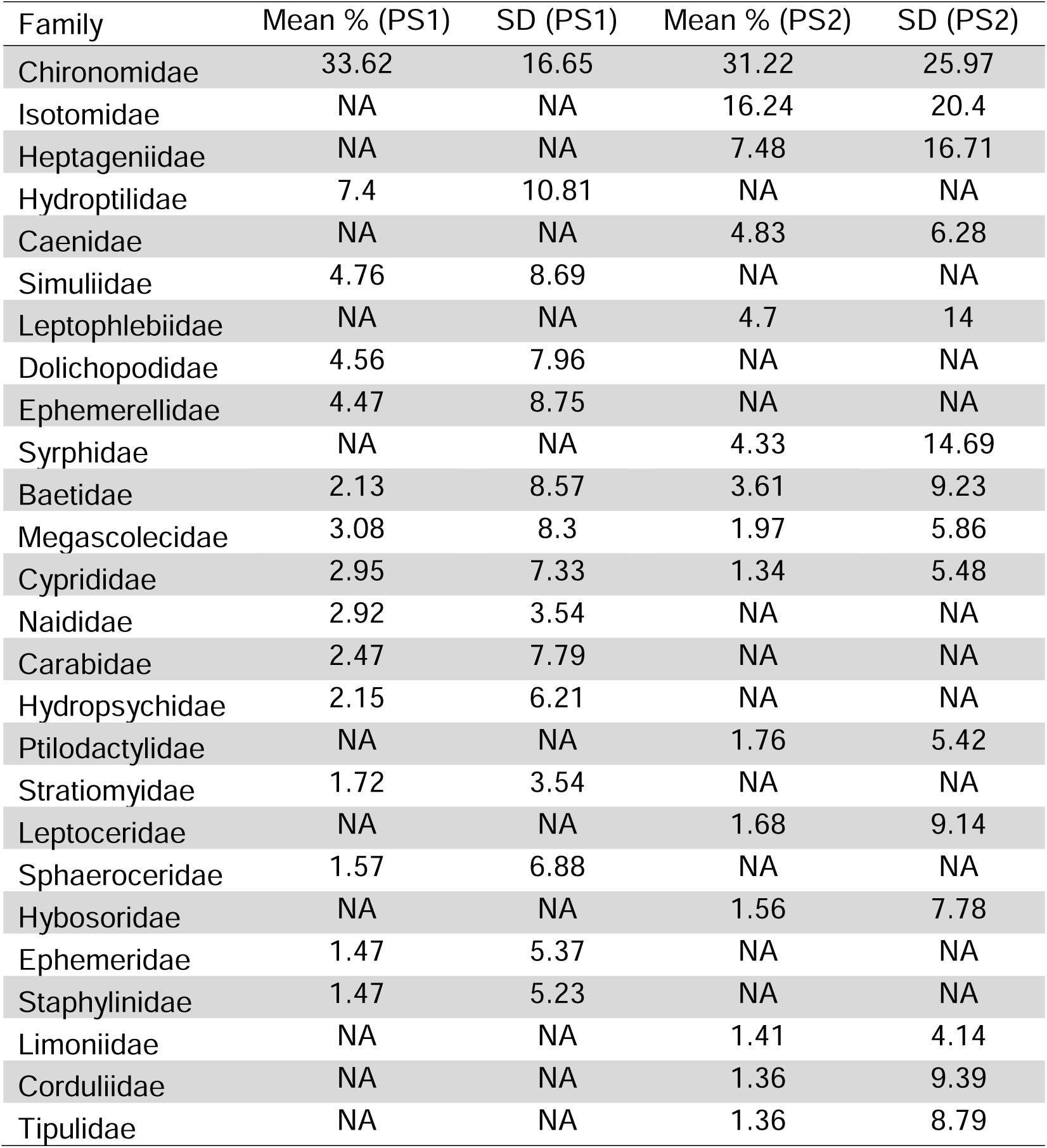
Diversity and Composition of Diet among Mormyrids of Balé Creek.

Beta diversity analysis using a comprehensive PERMANOVA framework with marginal (Type III) tests assessed the independent contributions of species identity, waveform type, sex, and body size to dietary variation. Species identity was the dominant predictor of diet composition, explaining ∼24% of variation in both primer sets (PS1: R² = 0.240, F_8,37_ = 1.69, *P* = 0.001; PS2: R² = 0.235, F_8,33_ = 1.49, *P* = 0.001). Body size explained a small but significant additional fraction of variation in PS1 (R² = 0.029, F_1,37_ = 1.63, *P* = 0.004), though this effect was not significant in PS2 (R² = 0.024, *P* = 0.158). Waveform type explained minimal unique variance after controlling for species (PS1: R² = 0.016, *P* = 0.720; PS2: R² = 0.011, *P* = 0.961), consistent with strong confounding between species identity and waveform—only *M. zanchlirostris* contained both BP and TP individuals at this site. Sex was not a significant predictor in either primer set (PS1: R² = 0.037, *P* = 0.321; PS2: R² = 0.044, *P* = 0.255).

Variance partitioning using adjusted R² values (Fig. 4D; Table S2), which account for the number of parameters in each predictor set, confirmed that ecological factors (species + waveform) explained substantially more unique variation (adjusted R² = 10.5–11.9%) than individual-level factors (sex + body size; adjusted R² = 1.2–1.8%), with minimal shared variance between these sets. Three-way partitioning (Table S3) further showed that species identity accounted for nearly all ecological variance (unique adjusted R² = 8.5–10.7%), while waveform contributed negligible unique variance when both species and individual factors were controlled (adjusted R² ≤ 0), reflecting the strong species–waveform confounding at this single site.

PCoA ordination revealed that non-*Paramormyrops* genera, particularly *Marcusenius moori* and *Mormyrops zanchlirostris*, occupied distinct dietary space from the *Paramormyrops* cluster, which showed substantial overlap (Fig. 4C). Dispersion tests indicated significant heterogeneity of variance among species (PS1: F = 35.9, *P* = 0.001; PS2: F = 10.6, *P* = 0.001), suggesting that the significant species effect in the PERMANOVA may be partly driven by differences in dietary variability rather than solely by centroid differences. In contrast, dispersion was homogeneous for waveform type and sex, supporting a valid interpretation of those PERMANOVA results (Table S4). Diet composition and diversity patterns were concordant between primer sets. Both primer sets showed consistent rankings of effect sizes, similar PCoA structure, and concordant prey family compositions, validating the robustness of our findings.

### Dietary Variation in Paramormyrops kingsleyae

#### Alpha Diversity

*Paramormyrops kingsleyae* are sexually dimorphic in size (males: 98 ± 23 mm; females: 87 ± 10 mm; *t*-test *p* = 0.023), and approximately 19% (22/119) of individuals harbored internal parasites, with significant variation in parasite prevalence between sites (Table S5). To establish the baseline structure of dietary variation, we first examined whether any of these factors affected dietary breadth before turning to compositional analyses.

*Paramormyrops kingsleyae*, like the mormyrids of Balé Creek, primarily consumed chironomids (>30% of diet composition), though the remaining prey consisted of highly diverse taxa including Baetidae, Carabidae, Naididae (Annelida), and Coenagrionidae (Odonata) (Fig. 5A). None of the five factors examined significantly affected dietary breadth. Shannon diversity did not differ between waveform types (Fig. 5B; two-way ANOVA: PS1 *F*_1,78_ = 0.003, *p* = 0.96; PS2 *F*_1,57_ = 1.422, *p* = 0.24) or regions (PS1 *F*_1,78_ = 2.078, *p* = 0.15; PS2 *F*_1,57_ = 2.406, *p* = 0.13), with a marginal Region × Waveform interaction in PS1 only (*F*_1,78_ = 3.455, *p* = 0.067). Males and females did not differ in dietary breadth when controlling for body size (ANCOVA: PS1 *F*_1,76_ = 6.826, *p* = 0.011; PS2 *F*_1,62_ = 0.059, *p* = 0.809). Body size showed a weak, non-significant negative correlation with Shannon diversity (PS1: *r* = −0.099, *p* = 0.336; PS2: *r* = −0.206, *p* = 0.075), and parasitism had no effect on dietary breadth after controlling for body size (PS1 ANCOVA *p* = 0.138; PS2 *p* = 0.929). The consistent absence of alpha diversity effects across all factors indicates that variation in *P. kingsleyae* diets is driven by differences in *which* prey taxa are consumed rather than *how many*, motivating the compositional analyses that follow.

**Figure 5.**
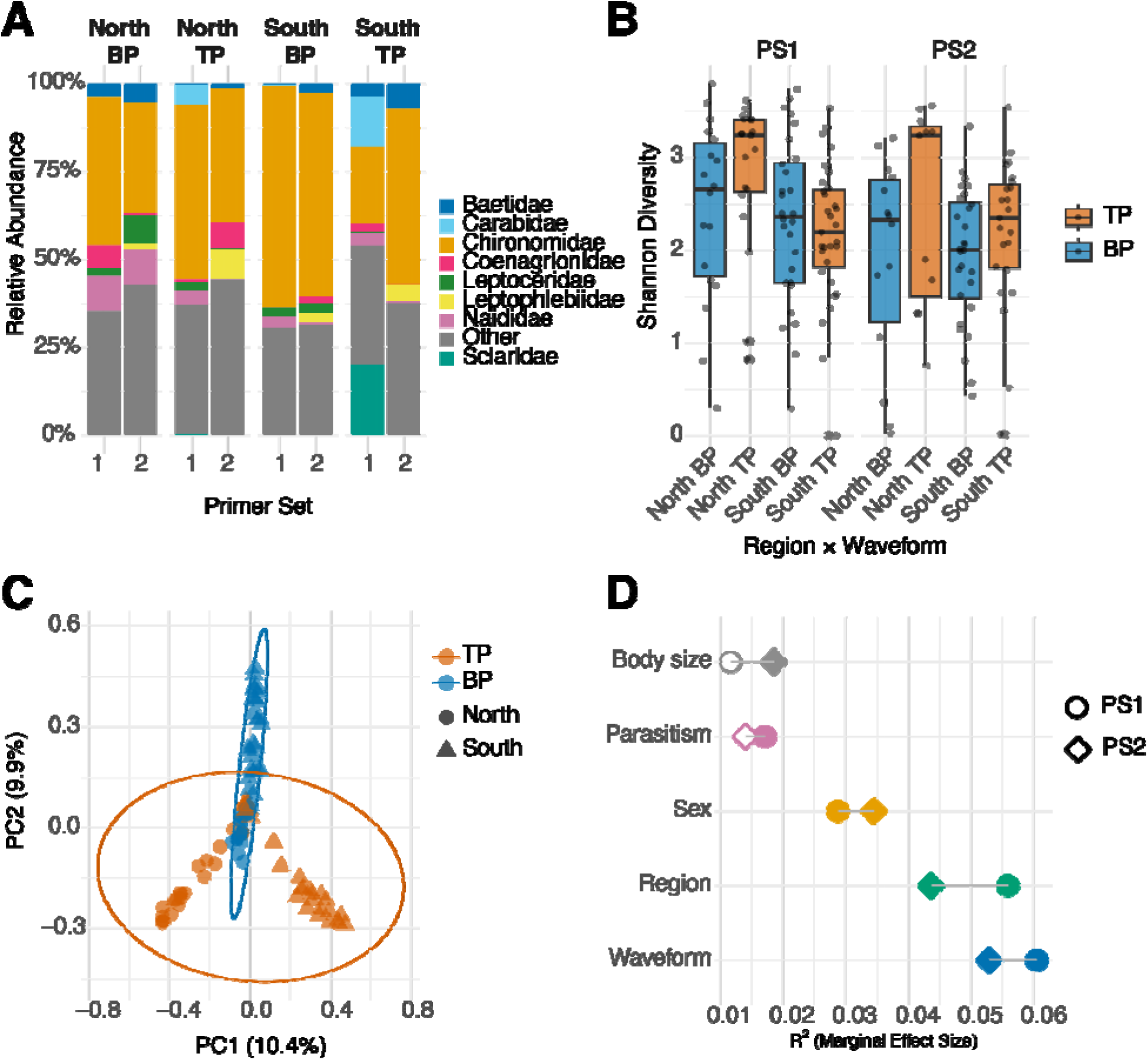
Dietary diversity across populations of *Paramormyrops kingsleyae*. A. Composition barplot of the top 20 prey taxa for primerset 1 and primerset 2 for each species. B. Mean Shannon diversity indices for each population, colored by waveform type. C. PCoA of Bray-Curtis dissimilarity among species for Primerset 1. 95% centroids for each species are shown, biphasic (BP) individuals are shown as triangles, triphasic (TP) individuals are shown as circles. D. Adjusted marginal effect sizes for waveform, body size, sex, and parasitism, and geographical region. Filled symbols are significant (p<0.05), empty symbols are not significant (p>0.05).

#### Beta Diversity: Relative Importance of Ecological and Individual Factors

To assess the relative importance of ecological and individual factors in shaping diet composition, we fit a comprehensive PERMANOVA model including waveform type, geographic region, sex, body size, and parasitism status as simultaneous predictors (Table S6). The model was run on 96 individuals (Primerset 1) and 76 individuals (Primerset 2) with complete data for all variables, including juveniles coded as a third level of sex.

Waveform type was the strongest independent predictor of diet composition, explaining 6.0% of variation after controlling for all other factors (Primerset 1: *R*^2^ = 0.060, *F*_1,89_ = 6.744, *p* = 0.001; Primerset 2: *R*^2^ = 0.053, *F*_1,69_ = 4.496, *p* = 0.001). Geographic region was the second-strongest predictor, explaining 5.6% (PS1: *R*^2^ = 0.056, *F*_1,89_ = 6.229, *p* = 0.001; PS2: *R*^2^ = 0.044, *F*_1,69_ = 3.713, *p* = 0.001). Sex explained 2.9–3.4% of variation (PS1: *R*^2^ = 0.029, *p* = 0.005; PS2: *R*^2^ = 0.034, *p* = 0.005), parasitism explained 1.4–1.7% (PS1: *R*^2^ = 0.017, *p* = 0.001; PS2: *R*^2^ = 0.014, *p* = 0.130), and body size explained 1.2–1.9% (PS1: *R*^2^ = 0.012, *p* = 0.097; PS2: *R*^2^ = 0.019, *p* = 0.014). These results were robust to the order in which terms entered the model with waveform showing particularly stable effect sizes across sequential analyses (Table S7).

Homogeneity of dispersion was confirmed for the focal factor of waveform type (betadisper: PS1 *F* = 2.019, *p* = 0.155; PS2 *F* = 0.019, *p* = 0.898), though significant dispersion heterogeneity was detected for region in Primerset 1 (*F* = 6.931, *p* = 0.012) and parasitism in both primer sets (PS1 *F* = 8.556, *p* = 0.005; PS2 *F* = 4.392, *p* = 0.049), suggesting these effects should be interpreted with caution.

Three-way variance partitioning (Fig. 5D; Table S9) confirmed that waveform and geography explained largely independent components of dietary variation. Pure waveform effects accounted for 5.4% (PS1) to 4.4% (PS2) of variance, pure geographic effects for 5.0% to 3.4%, with negligible shared variance between them (<0.5%). Individual factors (sex, body size, parasitism) collectively explained an additional 2.8% (PS1) to 2.2% (PS2) of unique variance. Ecological factors thus explained roughly four times more dietary variation than individual phenotypic factors (two-way partition: ecological 10.7% vs. individual 2.8% for PS1; 8.1% vs. 2.2% for PS2).

PCoA ordination of Bray-Curtis dissimilarity illustrated the modest but significant structuring detected by PERMANOVA (Fig. 5C). The first two axes explained 20.3% of total variation (PC1: 10.4%; PC2: 9.9%), with individuals distributed broadly across ordination space consistent with the extremely high overall dissimilarity among individuals (mean Bray-Curtis > 0.98). Northern BP and TP individuals overlapped extensively in ordination space, whereas southern BP and TP individuals showed greater separation from one another, suggesting that waveform-associated dietary divergence may be more pronounced in southern populations.

Geography contributed an additional structuring effect, with northern and southern individuals partially offset along PC1. Dispersion tests confirmed homogeneous variance for waveform type across both primer sets (betadisper: PS1 F = 2.019, p = 0.155; PS2 F = 0.019, p = 0.898) and for sex (PS1 F = 0.166, p = 0.844; PS2 F = 2.292, p = 0.103), supporting valid interpretation of these PERMANOVA results. However, significant dispersion heterogeneity was detected for parasitism in both primer sets (PS1 F = 8.556, p = 0.005; PS2 F = 4.392, p = 0.049) and for region in PS1 (F = 6.931, p = 0.012) but not PS2 (F = 1.630, p = 0.198), suggesting that the PERMANOVA results for these factors may partly reflect differences in dietary variability rather than solely centroid differences.

#### Waveform and Geographic Effects on Diet Composition

Having established that waveform type was the strongest predictor in the combined model, we examined the mechanisms underlying waveform and geographic dietary differences in detail. Overall, Bray-Curtis dissimilarity between BP and TP individuals was extremely high (PS1: 0.983; PS2: 0.987), and SIMPER analysis identified Chironomidae as the dominant family driving waveform differences, contributing 32.9% (PS1) to 29.6% (PS2) of total dissimilarity (Table S10). Beyond chironomids, the two primer sets revealed complementary sets of secondary prey families contributing to waveform differences: Primerset 1 highlighted Sciaridae (4.6%), Carabidae (4.0%), Naididae (2.9%), and Dolichopodidae (2.1%), while Primerset 2 identified Leptophlebiidae (4.2%), Coenagrionidae (2.7%), Ceratopogonidae (Diptera) (2.6%), and Baetidae (Ephemeroptera) (2.3%).

Dietary differences between BP and TP individuals were driven overwhelmingly by prey species turnover rather than differential abundance of shared prey (Table S12). Within the dominant Chironomidae, turnover accounted for 98.2% (PS1) to 98.3% (PS2) of Bray-Curtis dissimilarity, with only 17.8–18.0% of shared chironomid ASVs showing significant abundance differences between waveform types. BP and TP individuals shared 44.5% (PS1) to 44.7% (PS2) of chironomid ASVs (Jaccard similarity), indicating that more than half of the chironomid species consumed were unique to one waveform type or the other. At the whole-community level, BP and TP individuals shared 393 ASVs but differed in 206 BP-only and 249 TP-only ASVs (PS1; Jaccard = 0.463), with unique ASVs contributing 34.4% (BP) and 25.1% (TP) of total reads (Fig 6A). Among the top 15 ASVs driving waveform dissimilarity, the majority were chironomid species (Fig. 6B), but key non-chironomid taxa included a Sciaridae ASV, *Pterostichus* (Carabidae), *Platypalpus* (Diptera: Hybotidae), *Anisocentropus* (Trichoptera: Calamoceratidae), and *Nehalennia* (Coenagrionidae), reflecting the diverse array of invertebrate prey contributing to dietary divergence.

**Figure 6.**
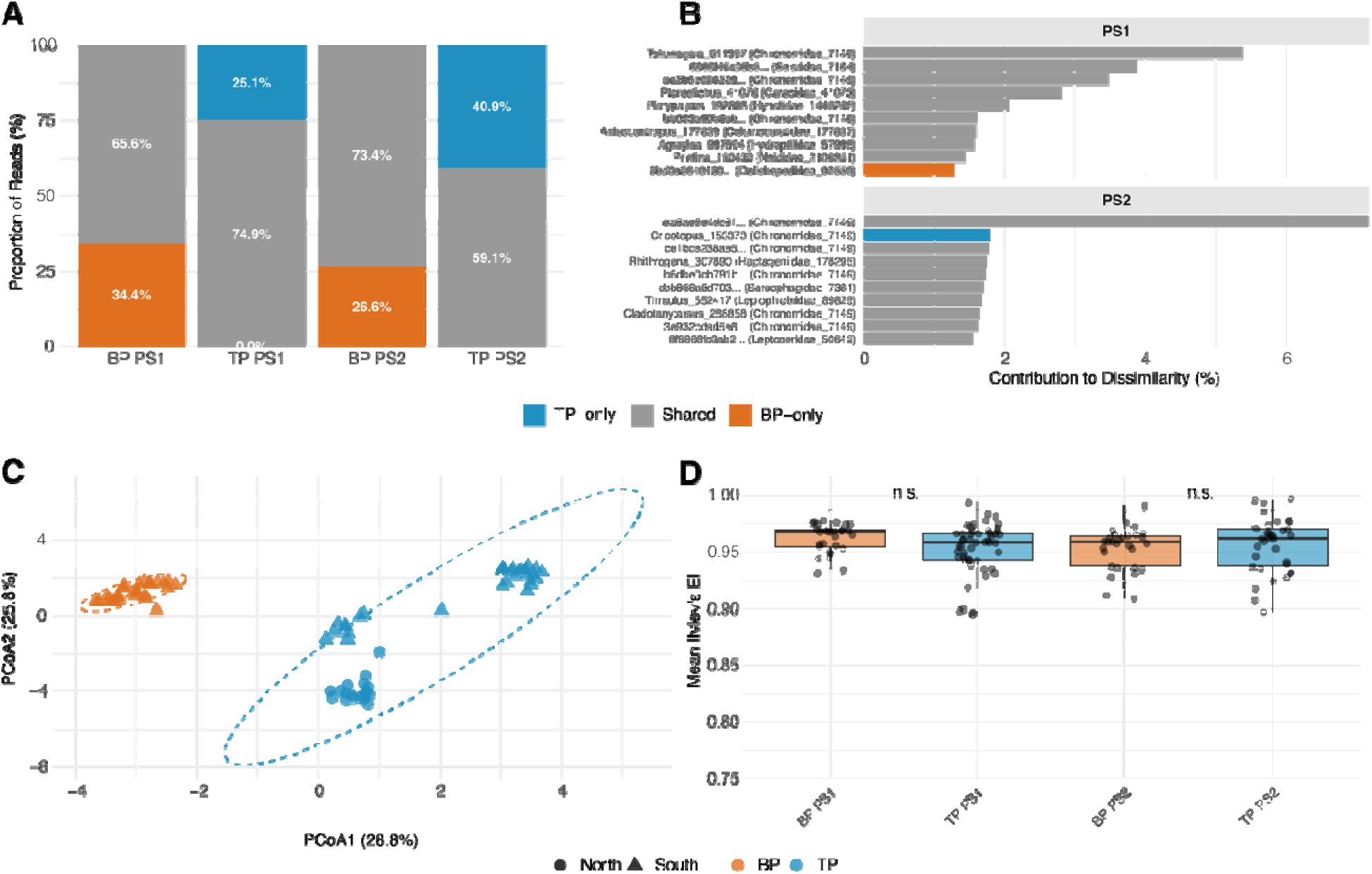
Dietary composition and prey selectivity differ between biphasic and triphasic *Paramormyrops kingsleyae* across both primer sets. *(A)* Stacked bar plots showing the proportional contribution of the three most abundant prey categories to gut contents of biphasic (BP, orange) and triphasic (TP, blue) *P. kingsleyae* as detected by primer set 1 (PS1) and primer set 2 (PS2). *(B)* SIMPER analysis identifying the prey ASVs contributing most to dietary dissimilarity between BP and TP individuals for PS1 (top) and PS2 (bottom); taxon labels indicate family assignment and database accession. *(C)* Principal coordinates analysis (PCoA) of Bray-Curtis dissimilarities among individual *P. kingsleyae* gut content samples, with color indicating waveform (BP orange; TP blue) and shape indicating region (Circles, North; Triangles South), dashed ellipses indicate 95% confidence regions. *(D)* Ivlev’s electivity index for individual *P. kingsleyae* relative to available prey, plotted by waveform type and primer set. Values near 1.0 indicate strong selectivity; both waveform types are highly selective but differ in which prey they preferentially consume (dashed line indicates the “equally selective, different preferences” threshold).

Geographic differences between northern and southern populations showed similar mechanistic patterns. Overall dissimilarity between regions was comparably high (PS1: 0.984; PS2: 0.989), and Chironomidae again dominated (PS1: 33.1%; PS2: 27.4%), followed by a similar suite of secondary families (Table S11). Within Chironomidae, turnover accounted for 99.0% (PS1) to 99.0% (PS2) of regional dissimilarity (Table S12) — slightly higher than for waveform differences — and Jaccard similarity was lower between regions (PS1: 0.355; PS2: 0.318) than between waveform types, indicating greater geographic than waveform-associated turnover in chironomid species. A higher proportion of shared chironomid ASVs showed significant regional abundance differences (PS1: 36.0%; PS2: 29.3%) compared to waveform-associated differences (17.8–18.0%), suggesting that while both factors operate primarily through species turnover, geographic effects additionally involve greater modulation of shared species abundances.

Critically, the near-zero shared variance between waveform and geography in the three-way partition (<0.5%) indicates that these represent distinct dietary signals: waveform-associated differences do not simply reflect the geographic structuring of prey communities.

#### Prey Selectivity

To distinguish whether waveform types consume different prey due to differential availability or differential selection, we calculated Ivlev’s electivity index for each individual based on environmental prey availability at each site (Fig. 6C). Waveform type was a strong predictor of selectivity patterns (Table S13; PS1: PERMANOVA *R*^2^ = 0.257, *F*_1,76_ = 38.276, *p* = 0.001; PS2: *R*^2^ = 0.344, *F*_1,60_ = 45.672, *p* = 0.001), explaining substantially more variation in selectivity profiles than in raw diet composition (where waveform explained 5–6%). Region also significantly predicted selectivity (PS1: *R*^2^ = 0.215; PS2: *R*^2^ = 0.185). The strength of the waveform effect from ∼6% in raw composition to ∼26–34% in selectivity space indicates that much of the compositional similarity between waveform types is attributable to shared environmental prey availability, and that the underlying prey preferences of BP and TP individuals diverge far more than their raw diets suggest.

However, overall selectivity strength did not differ between waveforms (Fig. 6D, Table S14): mean absolute electivity (|*E*|) was uniformly high for both BP (PS1: 0.963 ± 0.013; PS2: 0.954 ± 0.019) and TP (PS1: 0.953 ± 0.022; PS2: 0.956 ± 0.025) individuals (Wilcoxon PS1: *W* = 916, *p* = 0.053; PS2: *W* = 429, *p* = 0.384). This indicates that while BP and TP individuals actively select for different prey items, they are equally selective in their foraging — the difference lies in *what* they prefer, not *how strongly* they prefer it.

## Discussion

We used DNA metabarcoding of gut contents to test whether EOD waveform polymorphism in *Paramormyrops kingsleyae* is associated with dietary niche partitioning. We demonstrate first, at the community level, dietary divergence among mormyrid species tracks broad phylogenetic boundaries rather than EOD waveform type, with non-*Paramormyrops* genera occupying clearly distinct dietary niches from the closely-related *Paramormyrops* species flock. Second, within *P. kingsleyae*, waveform type is the single strongest predictor of diet composition, independently explaining 5–6% of dietary variation after controlling for geography, sex, body size, and parasitism. Third, and most compellingly, when we shift from asking what fish eat to what fish actively prefer relative to local prey availability, the waveform effect grows from ∼6% to 26–34% of variation in selectivity profiles—a fourfold amplification that reveals prey preferences diverging far more strongly than raw gut content compositions alone would suggest. Together, these results are consistent with the hypothesis that EOD waveform polymorphism facilitates dietary niche partitioning.

### Community-level dietary structure reflects phylogeny, not waveform

At Balé Creek, species identity explains approximately 24% of dietary variation—substantially more than any other factor. The PCoA ordination reveals a pattern consistent with phylogenetic signal in diet: non-*Paramormyrops* genera such as *Mormyrops zanchlirostris* and *Marcusenius moori* occupy clearly separated dietary space, while *Paramormyrops* species cluster together with considerable overlap. This broad structure echoes the finding of Arnegard et al. (2010) that trophic divergence among mormyrids is substantially slower than electric signal divergence—the *Paramormyrops* radiation has produced remarkable EOD diversity without comparable dietary differentiation at the genus level.

*Mormyrops zanchlirostris* stands out as a dietary specialist, exhibiting markedly lower Shannon diversity than all *Paramormyrops* species examined. This is consistent with the elongated, forcep-like jaw morphology of *Mormyrops*, which appears to reflect a specialization for extracting prey from crevices. The contrast with *Paramormyrops*, which shares more similar and relatively high dietary breadth across species, suggests that within the *Paramormyrops* clade, species diversification has proceeded along axes other than gross dietary specialization—consistent with the primacy of sexual selection on EODs proposed by Arnegard et al. (2010).

That waveform explained negligible unique variance at the Balé Creek community level (R² ≤ 0.016, p > 0.7) is unsurprising given that community-level analyses are dominated by the deep intergeneric differences. The Balé Creek community analysis thus provides important ecological context—showing that dietary differentiation is possible among mormyrids, and that the scale of that differentiation increases with phylogenetic distance—but is underpowered for detecting the subtler intraspecific waveform effects that the focused *P. kingsleyae* analysis is designed to detect.

### Waveform type predicts diet composition independently of geography

Our comprehensive PERMANOVA framework simultaneously estimated the independent contributions of waveform and region, and the results are unambiguous: both factors explain significant, largely non-overlapping fractions of dietary variation (waveform: 5.4–6.0%; geography: 3.4–5.6%; shared variance < 0.5%). The robustness of the waveform effect to term-ordering in sequential PERMANOVA tests further confirms that this is not an artifact of model specification.

The magnitude of the waveform effect (R² ≈ 0.05–0.06) is modest in absolute terms but is both statistically robust and ecologically interpretable. Small R² values are the norm in diet metabarcoding studies—individual dietary variation is extremely high, as reflected by the near-maximal mean Bray-Curtis dissimilarity among individuals (> 0.98). The biological question is not whether waveform explains most dietary variation (it does not, and we would not expect it to given opportunistic foraging, individual variation, and the many environmental factors shaping what any given fish encounters on any given day), but whether waveform explains a consistent, independent, and meaningful fraction of that variation. That we detect concordant waveform effects across two independent primer sets —that preferentially amplify different invertebrate communities—substantially strengthens confidence in this finding.

The mechanisms underlying waveform-associated dietary differences involve primarily prey species turnover rather than differential abundance of shared prey. Within Chironomidae alone—the dominant dietary family for both waveform types—turnover accounted for 98% of Bray-Curtis dissimilarity between waveform types, and fewer than half of chironomid ASVs were shared between BP and TP fish (Jaccard ≈ 0.45). This pattern has a clear biological interpretation: BP and TP fish are not consuming different amounts of the same chironomid species; they are consuming largely non-overlapping sets of chironomid species. At the resolution of individual ASVs, this pattern suggests that the dietary divergence operates at a finer taxonomic scale than is typically resolved by stable isotope or visual inspection of gut contents.

### Prey selectivity reveals divergent foraging preferences beyond raw diet composition

The prey selectivity analysis is the most powerful evidence that waveform-associated dietary differences reflect genuine preference rather than passive sampling of different prey environments. By comparing each individual fish’s gut contents against the invertebrate community available at its collection site, we convert absolute diet compositions into preference profiles that account for local prey availability. The waveform effect in this selectivity space (R² = 0.26–0.34) is roughly four to five times larger than the effect on raw diet composition (R² = 0.05–0.06). This amplification has a straightforward interpretation: a substantial portion of the raw dietary similarity between BP and TP fish is attributable to similar prey consumption because that prey is available, not because they prefer the same prey. When availability is controlled, the underlying preference divergence between waveform types becomes much clearer.

Importantly, this waveform effect on selectivity patterns cannot be explained by geographic differences in what prey is available because geographic region is also controlled as an independent predictor in the selectivity PERMANOVA and explains a separate fraction of variance. The near-zero shared variance between waveform and geography in the three-way partition confirms that these are distinct dietary signals: BP and TP fish actively prefer different prey even when foraging in environments with similar prey assemblages.

Equally notable is the finding that BP and TP individuals do not differ in overall selectivity intensity—both waveform types are equally selective (mean |E| ≈ 0.95–0.96), with high values indicating that neither waveform type feeds randomly relative to prey availability. The waveform effect lies entirely in the identity of preferred prey, not in the strength of selectivity. This result is inconsistent with a null model in which waveform types are ecologically equivalent generalists whose dietary differences arise from sampling different local prey communities. Instead, both waveform types appear to be comparably selective foragers targeting different prey items with equal discrimination.

### Why would waveform type predict prey choice? A candidate mechanism

The most surprising aspect of our findings is not that waveform-associated dietary differences exist, but that they exist at all given the physical similarity of BP and TP waveforms. BP and TP *P. kingsleyae* have essentially identical EOD power spectra — the frequency components present in each signal are indistinguishable — and the additional phase present in the TP waveform contributes negligibly to overall signal energy (Gallant et al., 2011; Picq et al., 2020). Yet, *P. kingsleyae* can reliably discriminate BP from TP waveforms behaviorally (Picq et al. 2020). Is it possible that EOD differences contribute to differences in prey perception as well?

Gottwald et al. (2018) demonstrated that the mormyrid *Gnathonemus petersii* can discriminate objects based on the ratio of amplitude modulation to waveform modulation imposed on the fish’s own EOD as it interacts with an object’s complex impedance — a ratio that is intrinsic to the object regardless of distance or size, analogous to color in vision.

Crucially, this discrimination relies not on peripheral receptor sensitivity alone, but on central subtraction of signals from A- and B-type mormyromast afferents (Gottwald et al., 2018; von der Emde & Bleckmann, 1992), with waveform extraction occurring at a higher processing level.

While BP and TP waveforms do not significantly differ in frequency content (Picq et al. 2020; Gallant et al. 2011), they differ in the *phase relationships* among those frequency components — the temporal structure of how they combine. When an EOD interacts with a capacitive object such as a prey item, that object imposes phase shifts on the returning signal that depend on its own RC time constant. Two waveforms with identical power spectra, but different phase structure, could produce measurably different distortions from the same capacitive object because the interaction between the outgoing signal’s temporal structure and the object’s impedance is phase-sensitive. If so, the relevant consequence would not be that BP fish can detect prey items TP fish cannot — rather, the *contrast among prey types* differing in capacitive properties might be enhanced differently for each waveform type. One waveform might provide sharper discrimination among prey items in one region of impedance space; the other might resolve a different cluster more crisply. This would produce exactly the pattern we observe: not wholesale avoidance of certain prey, but differential selection among prey categories that are both available and detected by both waveform types. This possibility becomes particularly compelling when the ecology of the dominant prey is considered.

Chironomid larvae are predominantly benthic burrowers, living concealed within sediment tubes or organic detritus, and many other prey recovered from guts — including mayfly and caddisfly larvae — similarly occupy cryptic microhabitats among substrate particles and leaf litter. For prey that are effectively invisible to visual or mechanical senses, electrolocation may be the primary detection modality, making even subtle waveform-mediated differences in “electric contrast” among buried or concealed prey potentially meaningful for foraging efficiency.

We emphasize that this mechanism is entirely speculative. Whether the mormyromast central subtraction circuit is calibrated to each fish’s individual EOD waveform — a prerequisite for phase structure to influence prey color perception — is unknown. Direct measurement of the electrical impedance properties of the prey taxa we identify in fish guts has not been done. And other explanations remain viable: BP and TP populations may differ in microhabitat use, diel activity, or other unmeasured behavioral characteristics that influence prey encounter independently of active electrolocation. What we can say with confidence is that the dietary differences are real, robust across two independent primer sets and two analytical frameworks, and cannot be attributed to differential prey availability. The mechanism linking waveform to foraging ecology is a compelling open question, and one that is perhaps more tractable than it appears: an experiment measuring the ‘electric colors’ of key prey taxa using BP and TP EOD waveforms as stimuli would be a direct test of whether the physical interaction differs in the way this framework predicts.

### Revisiting Arnegard et al. (2010) with enhanced resolution

Arnegard et al. (2010) concluded that sexual selection on electric signals was the primary driver of *Paramormyrops* diversification, based in part on the observation that EODs diverged 9–17 times faster than trophic ecology as measured by stable isotopes. Importantly, however, those authors also noted “smaller yet significant differences in all measured ecological traits” among closely related species, and explicitly acknowledged the limitations of stable isotope analysis for detecting fine-scale trophic divergence. Fifteen years later, we return to Balé Creek with a methodology capable of resolving those finer-scale differences.

Our findings do not contradict the central conclusion of Arnegard et al. (2010)—the dramatic EOD divergence across the *Paramormyrops* radiation clearly outpaces gross dietary differentiation, and the community-level analysis confirms that *Paramormyrops* species overlap substantially in dietary space. However, by focusing on the intraspecific EOD polymorphism in *P. kingsleyae* and using prey selectivity to distinguish preference from availability, we reveal a layer of dietary divergence associated with waveform type that is precisely the kind of “small but significant ecological difference” the isotope approach was too coarse to detect. The electric color hypothesis predicts dietary divergence at the level of specific prey taxa within shared prey categories—exactly the level at which ASV-resolution metabarcoding operates and at which stable isotopes are essentially blind.

Our study also advances beyond the work of Amen et al. (2024) on *Campylomormyrus*, which demonstrated waveform-correlated dietary differences in mormyrids using a DNA-based approach, but could not determine whether those differences reflected active preference or differential prey availability. By pairing gut content metabarcoding with environmental prey sampling at the same sites, we directly address the availability question that Amen et al. (2024) identified as the key outstanding limitation. The amplification of the waveform effect from raw composition to selectivity space provides the empirical answer: dietary differences between waveform types in *P. kingsleyae* substantially exceed what would be expected from differences in prey availability alone. Additionally, because *P. kingsleyae* is morphologically conserved across waveform types—lacking the dramatic snout morphology differences of *Campylomormyrus*—any dietary divergence we detect is more parsimoniously attributed to sensory differences in electrolocation than to mechanical constraints on prey capture.

### Methodological contributions and cross-primer validation

The use of two independent COI primer sets as internal validation is a key methodological contribution of this study. The two primer sets amplify partially overlapping but compositionally distinct prey communities: PS1 shows stronger amplification of certain Diptera and microcaddisflies (Trichoptera: Hydroptilidae), while PS2 more efficiently detects mayfly diversity (Ephemeroptera). Despite these compositional differences, both primer sets reveal concordant biological patterns—the same predictors are significant, with similar effect sizes and directionally consistent prey family contributions to dietary differences. This cross-primer concordance substantially elevates confidence in our conclusions beyond what either primer set alone could provide. We recommend multi-primer approaches as a general best practice in dietary metabarcoding studies, particularly when the goal is detecting subtle compositional differences rather than broadly characterizing diet.

The local barcode reference database, generated from 429 COI barcodes of macroinvertebrates collected at the study sites, proved essential for accurate taxonomic assignment of ecologically important prey families. For PS2, local barcodes provided the sole reference coverage for Heptageniidae and contributed 88% of Chironomidae assignments, which is particularly critical given that Chironomidae are the dominant prey family and the primary taxon driving waveform-associated dietary differences. The chronic underrepresentation of Central African freshwater invertebrate diversity in global barcode databases (BOLD, NCBI) means that studies relying exclusively on public references would substantially misclassify or fail to assign a large fraction of locally important prey taxa. Investment in regional reference barcode generation is thus not merely a technical refinement but a prerequisite for accurate dietary characterization in biodiverse tropical systems.

The discovery that host DNA removal timing critically affects cross-primer concordance deserves specific attention. When Chordata sequences were removed after rarefaction rather than before, host DNA contamination—which varied among samples—biased prey read counts in ways that generated apparent discordances between primer sets. Filtering host sequences before rarefaction allows proper standardization of prey read depth across samples regardless of contamination level, recovering concordant patterns. This finding has broad implications for diet metabarcoding study design: studies that rarefy before removing host contamination may introduce systematic biases that undermine cross-validation and inflate apparent inconsistencies among primer sets or methodological replicates.

### Limitations and future directions

A notable secondary finding of this study is that 19% of *P. kingsleyae* individuals harbored internal parasites, identified morphologically and molecularly as larvae of *Eustrongylides* sp. (Dioctophymatidae). The identification itself yielded an unexpected result: nanopore assembly of the COI locus produced an ∼11 kb contig rather than the anticipated ∼650 bp barcode fragment, consistent with the fragmented mitochondrial architecture recently described in the related dioctophymatid *Dioctophyme renale* (Macchiaroli et al., 2025), in which each protein-coding gene occupies its own circular minichromosome flanked by an extensive non-coding region (Macchiaroli et al., 2025). The life cycle of *Eustrongylides* requires piscivorous birds as definitive hosts (Lichtenfels & Stroup, 1985; Honcharov et al., 2022), raising the untested possibility that fish-eating birds prey on *P. kingsleyae*. Parasitism rates varied dramatically across sites, from 0% at Balé Creek to 54% at Ipassa Creek (Table S5), likely reflecting local differences in oligochaete intermediate host abundance. These observations fall outside the central questions of this study but merit dedicated follow-up.

The present study also has limitations that future work should address. The geographic confound between waveform type and sampling locality cannot be fully eliminated with observational data from this system. Our variance partitioning approach provides strong analytical evidence that waveform effects are independent of geography, but we cannot rule out the possibility that unmeasured site-level variables correlated with waveform type contribute to the observed dietary differences. Future work using experimental manipulation of foraging behavior in controlled environments—providing fish with standardized prey assemblages and recording EOD-mediated detection behaviors—would provide the causal evidence that field studies inherently cannot.

The temporal snapshot nature of gut content sampling is a further limitation: diet composition captured at the moment of collection reflects what each fish encountered and chose to eat in the preceding hours, not its long-term dietary preferences. Seasonal variation in prey availability and fish foraging behavior could introduce noise that masks or inflates dietary differences between waveform types. Although our large sample size (n = 119 *P. kingsleyae* individuals) and the cross-primer concordance of waveform effects suggest that the patterns we detect are robust to this individual-level variation, studies spanning multiple seasons would strengthen inferences about whether waveform-diet associations are consistent features of the system.

The mechanisms connecting EOD waveform to prey detection remain entirely unknown. We have outlined a speculative candidate involving differential phase interactions with prey impedance, but this has not been tested directly. Characterizing the complex impedance of key prey families identified in fish guts — and measuring the distortion signatures those objects produce when illuminated with BP versus TP waveforms — would be a tractable next step.

### Conclusions

We have demonstrated that EOD waveform type is the strongest predictor of dietary composition in *Paramormyrops kingsleyae*, independently of geography, sex, body size, and parasitism status. The waveform effect is concordant across two independent primer sets, is driven by prey species turnover rather than differential abundance of shared prey, and is amplified fourfold when prey availability is controlled—indicating that BP and TP fish actively prefer different prey rather than passively encountering different prey communities. These findings suggest that natural selection on foraging efficiency may contribute alongside sexual selection to maintaining EOD waveform polymorphism in *P. kingsleyae*. More broadly, our results illustrate that the “small but significant” ecological differences noted by Arnegard et al. (2010) in the *Paramormyrops* radiation are real and ecologically meaningful, but were invisible to the isotope-based approaches available fifteen years ago. As DNA metabarcoding techniques continue to improve in taxonomic resolution and accessibility, and as local reference databases for underrepresented biogeographic regions expand, we anticipate that fine-scale dietary niche partitioning of this kind will prove to be a widespread feature of rapidly diversifying groups where signal evolution and sensory ecology are intimately linked.

## Acknowledgements

Permits to collect fishes in Gabon and export them for this study were granted by l’Institut de Recherche en Ecologie Tropicale, l’Institut de Recherches Agronomiques et Forestières, and the Centre National de la Recherche Scientifique et Technologique. We are grateful for the valuable assistance and logistical support we received from J. D. Mbega and Jean Hervé Mve Beh, as well as students working in these institutions.

## Author contributions

Conceptualization: JRG, SP. Funding Acquisition: JRG. Supervision: JRG. Methodology: JRG, SP, LK, NNA, FN, HKM. Investigation: SP, RG, EP, RB, MEB, ECK, RCS. Formal Analysis: SP, RG, JRG. Visualization: JRG. Data Curation: JRG, SP. Writing – Original Draft: JRG. Writing – Review & Editing: JRG, SP, RG, ECK, RCS, EP, MEB, LK, NNA, FN, HKM.

## Data Accessibility Statement

Raw amplicon sequence reads for this project are deposited in the NCBI Sequence Read Archive (BioProject: PRJNA1427285). Locally-generated COI barcodes from macroinvertebrates collected at study sites are deposited in the NCBI Nucleotide Database as PZ248292 - PZ248701. Various sequences obtained from Eustrongylides sp. parasites were generated: ITS region (Genbank PZ255420), COX1 (Genbank PZ259218). The full mitochondrial assembly is provided as Supplemental File 4. Sample metadata, including georeferenced collection localities, are provided in the Supplemental Data File 1. All analysis scripts are available on GitHub (https://github.com/msuefishlab/gut_contents_2025).

## Benefit-Sharing Statement

This research was conducted in compliance with Gabonese national law and applicable provisions of the Convention on Biological Diversity and Nagoya Protocol. A research collaboration was developed with scientists from Gabon’s Centre National de la Recherche Scientifique et Technologique (CENAREST); all Gabonese collaborators are included as co-authors on this publication. Results of this research have been shared with collaborating institutions in Gabon and are being made available to the broader scientific community through open-access data deposition as described above. The research contributes to foundational knowledge of Gabonese freshwater biodiversity, including the first COI barcode library for macroinvertebrate communities at multiple Gabonese river systems, which will serve as a long-term resource for future ecological and biodiversity research in the region.

