## Supplemental Figures for "Electric signal polymorphism predicts dietary niche partitioning in a weakly electric fish"

***Electronic Supplement for:***

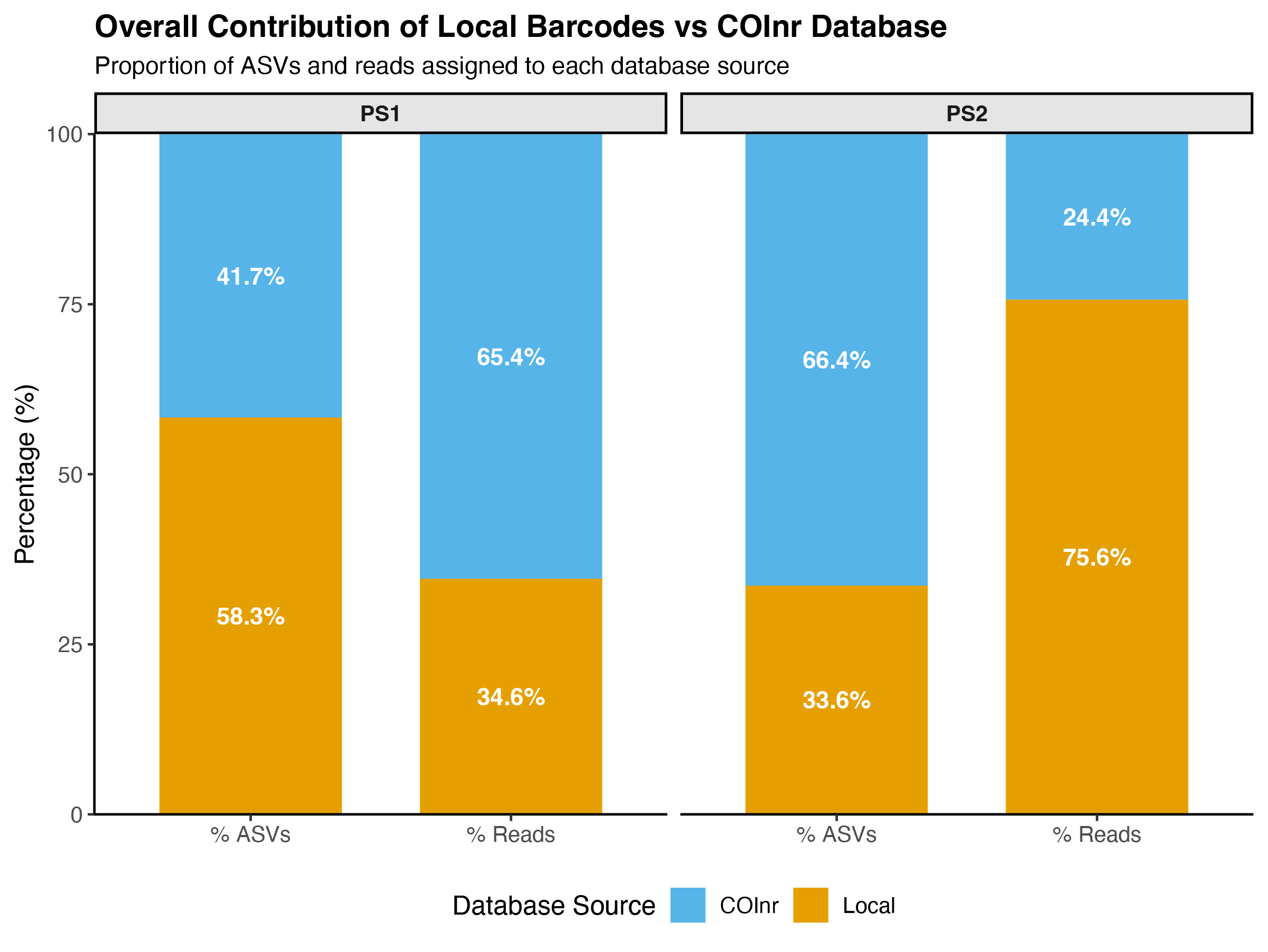

**Figure S1: Relative Contributions of COInr and Local Barcodes Sequenced from Collecting Sites.**

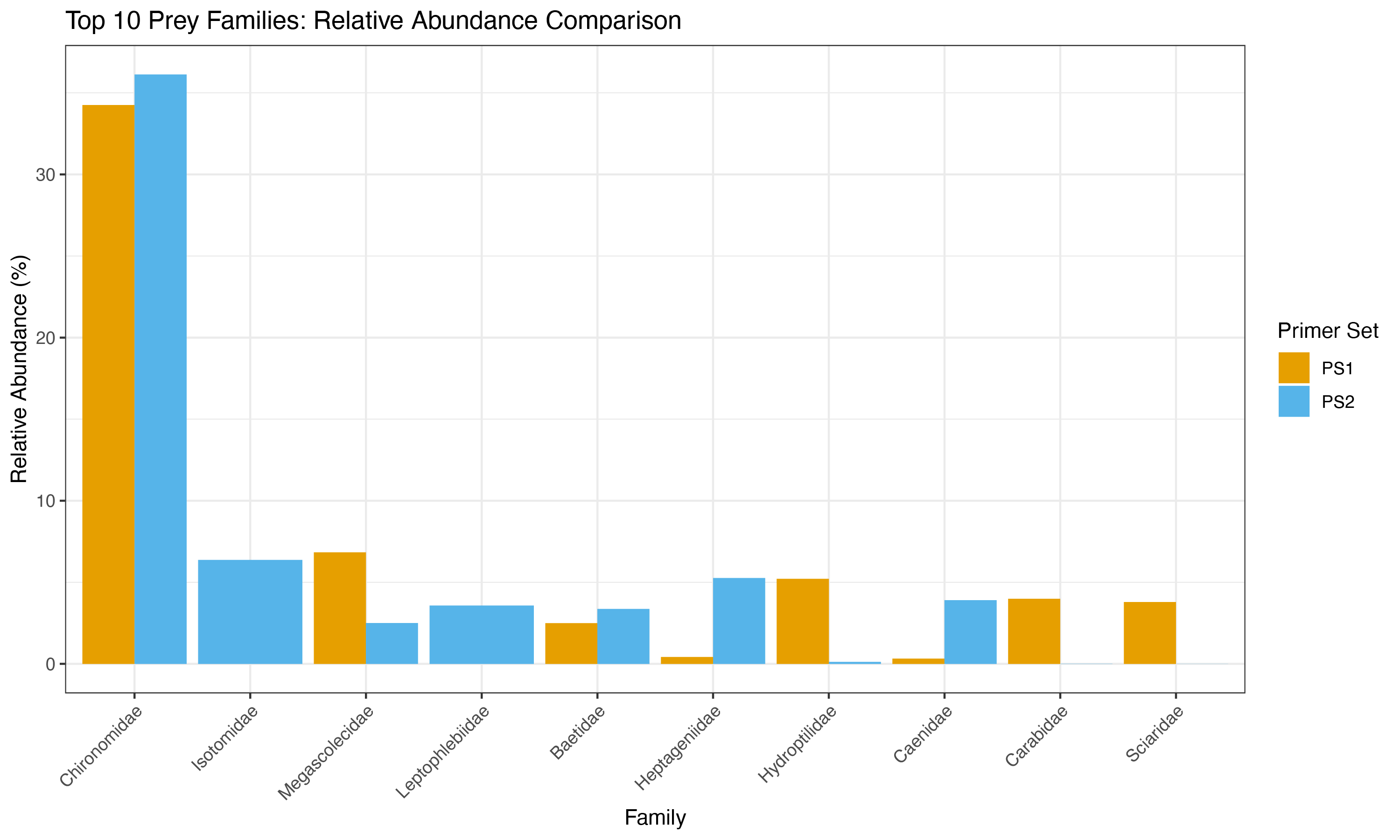

**Figure S2: Overlapping but compositionally distinct prey communities.** Comparison of prey communities identified using Primerset 1 (PS1) and Primerset 2 (PS2). While both primersets capture broadly similar ecological patterns, they reveal compositionally distinct subsets of the total prey diversity.

| Species | PS1 (n) | PS2 (n) |
| --- | --- | --- |
| *Marcusenius moori* | 9 | 9 |
| *Mormyrops zanchlirostris* | 7 | 6 |
| *Paramormyrops hopkinsi* | 1 | 1 |
| *Paramormyrops kingsleyae* | 9 | 6 |
| *Paramormyrops spp. MAG* | 2 | 2 |
| *Paramormyrops spp. SZA* | 11 | 10 |
| *Paramormyrops spp. TEN* | 1 | 2 |
| *Paramormyrops spp. TEU* | 0 | 0 |
| *Paramormyrops spp. VAD* | 1 | 1 |
| *Petrocephalus simus* | 9 | 9 |
| Total | **50** | **46** |

**Table S1. Sample Size Summary by Species and Primerset (Bale Creek).** *Data represents samples remaining after complete-case filtering. Species with n < 3 are included for transparency but were excluded from downstream PERMANOVA stability.*

| Primerset | Factor | Df | R2 | F | p-value |
| --- | --- | --- | --- | --- | --- |
| PS1 | Species | 8 | 0.2400 | 1.6937 | 0.001 *** |
|  | Waveform Type | 1 | 0.0158 | 0.8921 | 0.720 |
|  | Sex | 2 | 0.0371 | 1.0485 | 0.321 |
|  | Body size | 1 | 0.0288 | 1.6283 | 0.004 ** |
| PS2 | Species | 8 | 0.2348 | 1.4920 | 0.001 *** |
|  | Waveform Type | 1 | 0.0114 | 0.5777 | 0.961 |
|  | Sex | 2 | 0.0436 | 1.1070 | 0.255 |
|  | Body size | 1 | 0.0245 | 1.2441 | 0.158 |

**Table S2. Marginal PERMANOVA Results (Community-Level).** *Marginal (Type III) tests assessing independent contributions of ecological and individual factors.*

| Component | PS1 (Adj. R2) | PS2 (Adj. R2) |
| --- | --- | --- |
| Pure Ecological [a] (Species + Waveform) | 0.1188 | 0.1054 |
| Pure Individual [b] (Sex + Body size) | 0.0183 | 0.0120 |
| Shared [c] | -0.0048 | -0.0027 |
| Residual [d] | 0.8678 | 0.8853 |

**Table S3. Community Variance Partitioning (Ecological vs. Individual).** *Values represent Adjusted R-squared components calculated via dbRDA.* Note: Negative Adjusted R-squared values occur when the explanatory power of a factor is less than expected by random chance, indicating the factor explains negligible variance beyond stochastic noise.

| Factor | PS1 F | PS1 p | PS2 F | PS2 p | Note |
| --- | --- | --- | --- | --- | --- |
| Species | 35.935 | 0.001 | 10.588 | 0.001 | **Heterogeneous dispersion** |
| Waveform | 2.978 | 0.103 | 1.710 | 0.225 | Homogeneous |
| Sex | 0.712 | 0.514 | 0.050 | 0.944 | Homogeneous |

**Table S4. Dispersion Test Results (Community-Level).** *Testing the homogeneity of multivariate dispersion (PERMDISP).*

| Site | Mormyrids | *P. kingsleyae* | Parasite | Percent Parasitized |
| --- | --- | --- | --- | --- |
| Bale Creek | 59 | 9 | 0 | 0.00% |
| Bambomo Creek | 33 | 33 | 4 | 12.12% |
| Bengue Creek | 2 | 0 | 0 | 0.00% |
| Emon Creek | 1 | 1 | 0 | 0.00% |
| Iboundji Creek | 27 | 27 | 1 | 3.70% |
| Louetsi River | 15 | 0 | 0 | 0.00% |
| Mengono Creek | 25 | 25 | 4 | 16.00% |
| Ipassa Creek | 24 | 24 | 13 | 54.17% |
| Total | **186** | **119** | **22** | **18.49%** |

**Table S5. Summary of parasitism rates at different collecting sites throughout Gabon.**

| Primerset | Term | Df | R2 | F | P-value |
| --- | --- | --- | --- | --- | --- |
| PS1 | Region | 1 | 0.0558 | 6.2287 | 0.001 *** |
|  | Waveform Type | 1 | 0.0605 | 6.7439 | 0.001 *** |
|  | Sex | 2 | 0.0288 | 1.6058 | 0.005 ** |
|  | Body size | 1 | 0.0117 | 1.3018 | 0.097 . |
|  | Parasite status | 1 | 0.0172 | 1.9132 | 0.001 *** |
| PS2 | Region | 1 | 0.0436 | 3.7128 | 0.001 *** |
|  | Waveform Type | 1 | 0.0528 | 4.4961 | 0.001 *** |
|  | Sex | 2 | 0.0345 | 1.4670 | 0.003 ** |
|  | Body size | 1 | 0.0186 | 1.5803 | 0.011 * |
|  | Parasite status | 1 | 0.0141 | 1.1970 | 0.134 |

**Table S6. Comprehensive Marginal PERMANOVA (*P. kingsleyae*).** *Additive model assessing independent dietary predictors across all sampled regions.*

| Primerset | Term | Marginal R2 | Geo_First_R2 | Ind_First_R2 |
| --- | --- | --- | --- | --- |
| PS1 | Region | 0.0558 | 0.0722 | 0.0601 |
|  | Waveform Type | 0.0605 | 0.0666 | 0.0605 |
|  | Sex | 0.0288 | 0.0343 | 0.0382 |
| PS2 | Region | 0.0436 | 0.0585 | 0.0471 |
|  | Waveform Type | 0.0528 | 0.0624 | 0.0528 |
|  | Sex | 0.0345 | 0.0356 | 0.0461 |

**Table S7. Sequential PERMANOVA Sensitivity Analysis.** *Comparing R2 values based on term entry order to assess robustness against confounding.*

| Factor | PS1 F | PS1 p | PS2 F | PS2 p | Diagnostics |
| --- | --- | --- | --- | --- | --- |
| Waveform Type | 2.019 | 0.143 | 0.019 | 0.887 | Assumption Met |
| Region | 6.931 | 0.011 | 1.630 | 0.194 | Heterogeneous (PS1) |
| Sex | 0.166 | 0.852 | 2.292 | 0.107 | Assumption Met |
| Parasite Status | 8.556 | 0.005 | 4.392 | 0.049 | Heterogeneous |

**Table S8. Dispersion Diagnostics (*P. kingsleyae*).** *Verification of PERMANOVA assumptions via multivariate dispersion tests.*

| Component | PS1 (Adj. R2) | PS2 (Adj. R2) |
| --- | --- | --- |
| Pure Region | 0.0495 | 0.0342 |
| Pure Waveform | 0.0544 | 0.0440 |
| Pure Individual | 0.0280 | 0.0218 |
| Shared Region + Waveform | 0.0033 | 0.0025 |
| Shared Region + Individual | 0.0036 | 0.0070 |
| Shared Waveform + Individual | 0.0144 | 0.0131 |
| Shared All Three | -0.0049 | -0.0040 |

**Table S9. Variance Partitioning Components (*P. kingsleyae*).** *Adjusted R-squared values for three-way partitioning of dietary variance.*

| Primerset | Family | Contribution (%) | N ASVs |
| --- | --- | --- | --- |
| PS1 | Chironomidae | 32.86 | 39 |
|  | Sciaridae | 4.58 | 2 |
|  | Carabidae | 3.96 | 3 |
| PS2 | Chironomidae | 29.56 | 25 |
|  | Leptophlebiidae | 4.20 | 4 |
|  | Coenagrionidae | 2.65 | 3 |

**Table S10. SIMPER Analysis of Waveform Differences (BP vs. TP).** *Top prey families driving dissimilarity in P. kingsleyae.*

| Primerset | Family | Contribution (%) | N ASVs |
| --- | --- | --- | --- |
| PS1 | Chironomidae | 33.14 | 38 |
|  | Sciaridae | 3.83 | 2 |
|  | Carabidae | 3.80 | 3 |
| PS2 | Chironomidae | 27.40 | 28 |
|  | Leptophlebiidae | 3.59 | 4 |
|  | Coenagrionidae | 3.28 | 3 |

**Table S11. SIMPER Analysis of Regional Differences (North vs. South).**

| Comparison | Primerset | BC_Total | BC_Turnover_Pct | BC_Abundance_Pct | Dominant Mechanism |
| --- | --- | --- | --- | --- | --- |
| Waveform | PS1 | 0.981 | 98.24% | 1.76% | Turnover |
| Waveform | PS2 | 0.979 | 98.27% | 1.73% | Turnover |
| Region | PS1 | 0.979 | 99.01% | 0.99% | Turnover |
| Region | PS2 | 0.980 | 98.99% | 1.01% | Turnover |

**Table S12. Turnover vs. Abundance.** *Detailed partitioning of dissimilarity for Chironomidae, the primary driver of dietary variance.*

| Primerset | Factor | Df | R2 | F | Pr(>F) |
| --- | --- | --- | --- | --- | --- |
| PS1 | Waveform | 1 | 0.2573 | 38.276 | 0.001 *** |
|  | Region | 1 | 0.2154 | 32.040 | 0.001 *** |
| PS2 | Waveform | 1 | 0.3439 | 45.672 | 0.001 *** |
|  | Region | 1 | 0.1847 | 24.528 | 0.001 *** |

**Table S13. Individual Prey Selectivity PERMANOVA.** *Model: selectivity_dist ~ Waveform + Region. Results reflect marginal effects.*

| Primerset | W-statistic | p-value | Interpretation |
| --- | --- | --- | --- |
| PS1 | 916 | 0.0527 | Equally selective (n.s.) |
| PS2 | 429 | 0.3838 | Equally selective (n.s.) |

**Table S14. Mean Absolute Selectivity Comparison (Wilcoxon Tests).** *Testing for differences in "choosiness" (Mean |Ivlev's E|) between BP and TP waveforms.*
